# New Features Surrounding the Cdc42-Ste20 Module that Regulates MAP Kinase Signaling in Yeast

**DOI:** 10.1101/2023.02.28.530426

**Authors:** Beatriz González, Mahnoosh Mirzaei, Sukanya Basu, Aditi Prabhakar, Paul J. Cullen

## Abstract

Mitogen-activated protein kinase (MAPK) pathways regulate multiple cellular responses, including the response to stress and cell differentiation, and are highly conserved across eukaryotes from yeast to humans. In yeast, the canonical activation of several MAPK pathways includes the interaction of the small GTPase Cdc42p with the p21-activated kinase (PAK) Ste20p. We recently found that the active conformation of Cdc42p is regulated by turnover, which impacts the activity of the pathway that regulates filamentous growth (fMAPK). Here, we show that Ste20p is turned over by the 26S proteasome. Ste20p was stabilized when bound to Cdc42p, presumably to sustain MAPK pathway signaling. Ste20p is a major conduit by which signals flow through the fMAPK pathway; however, by genetic approaches we also identified a Ste20p-independent branch of the fMAPK pathway. Ste20p-dependent signaling required the 14-3-3 proteins, Bmh1p and Bmh2p, while Ste20p-independent signaling required the fMAPK pathway adaptor and Cdc42p-interacting protein, Bem4p. Ste20p-independent signaling was inhibited by one of the GTPase-activating proteins for Cdc42p in the fMAPK pathway, Rga1p, which also dampened basal but not active fMAPK pathway activity. Finally, the polarity adaptor and Cdc42p-interacting protein, Bem1p, which also regulates the fMAPK pathway, interacts with the tetra-span protein Sho1p, connecting a sensor at the plasma membrane to a protein that regulates the GTPase module. Collectively, these data reveal new regulatory features surrounding a Rho-PAK module that may extend to other pathways that control cell differentiation.

## INTRODUCTION

Mitogen-activated protein kinase (MAPK) pathways are evolutionary conserved modules that regulate a multiplicity of cellular responses including the response to stress (like osmotic stress), as well as cell proliferation, differentiation, and survival in eukaryotic cells (Braicu et al., 2019; Raman et al., 2007). In response to extracellular stimuli, MAPK pathways are activated by a diverse collection of receptors and sensors that typically function at the cell surface. Once activated, sensor proteins control the activity of G-proteins (heterotrimeric and monomeric) that govern the activity of effector kinases, commonly members of the p21-activated (PAK) family (Rane and Minden, 2014). The effector module of G-proteins and PAK kinases includes three kinases in a tandem series: MAP kinase kinase kinase (MAPKKK), MAP kinase kinase (MAPKK), and MAP kinase (Kolch, 2005; Morrison, 2012). One of the main functions of MAP kinases is the phosphorylation and activation of transcription factors that alter gene expression to mount a biological response. MAPK pathways can be induced by multiple stimuli, and accordingly, the core module can be ‘shared’ by different pathways to permit specific responses in different settings. Cross-talk between pathways occurs in normal settings (Fey et al., 2012; Kim and Rose, 2022), but when mis-regulated can alter normal cellular responses and lead to diseases including cancer (Braicu et al., 2019; Fey et al., 2012; Van Drogen et al., 2020). Understanding how pathways are regulated and induce pathway-specific signals is critical to understanding pathway mis-regulation and disease in higher organisms.

One model to study MAPK pathways is the budding yeast *Saccharomyces cerevisiae*. Three of the five yeast MAPK pathways are regulated by the Rho GTPase Cdc42p and PAK kinase Ste20p [(Saito, 2010; Schwartz and Madhani, 2004) filamentous growth, mating, and HOG]. The MAPK pathway that controls filamentous growth (fMAPK) induces cell differentiation in response to nutrient (nitrogen or carbon) limitation (Gimeno et al., 1992; Roberts and Fink, 1994). Filamentous growth is an underlying cause of virulence in plant and animal pathogens (Min et al., 2020; Mitchell, 1998). In *S. cerevisiae,* the fMAPK pathway is regulated by the mucin-type glycoprotein Msb2p (Cullen et al. 2004), the tetra-span osmosensor Sho1p (Adhikari et al., 2015; Yamamoto et al., 2010), and an integral membrane protein called Opy2p (Karunanithi and Cullen, 2012; Tatebayashi et al., 2015), whose main function is the plasma-membrane recruitment of the MAPKKK Ste11p (Truckses et al., 2006). Once activated, Msb2p and Sho1p regulate the Rho-type GTPase Cdc42p, which is a master regulator of signaling and cell polarity in eukaryotes. Cdc42p regulates the fMAPK pathway by proteins that converge on and regulate the guanine nucleotide exchange factor (GEF) for Cdc42p, called Cdc24p. Cdc42p activity is also controlled by GTPase activating proteins, one being Rga1p, the primary GAP for Cdc42p that regulates the fMAPK pathway (Smith et al., 2002). Among the proteins that regulate the Cdc42p module are the fMAPK scaffold Bem4p (Pitoniak et al., 2015), the Ras-type GTPase Rsr1p (Basu et al., 2020), which mainly controls bud-site selection (Park et al., 2002), and the polarity scaffold Bem1p (Basu et al., 2020), which regulates Cdc42p function during polarity establishment and signaling (Irazoqui et al., 2003; Leeuw et al., 1995; Lyons et al., 1996; Winters and Pryciak, 2005).

The main target of the Cdc42p-Ste20p module is the MAPKKK Ste11p, which functions to phosphorylate and activate the MAPKK Ste7p, which in turn phosphorylates the MAPK Kss1p (Leberer et al., 1997; Leeuw et al., 1998; Nagiec et al., 2015; Peter et al., 1996). Phosphorylation of Kss1p causes activation of the protein, and relief of its inhibitory functions (Cook et al., 1997), which leads to the induction of transcription factors including Ste12p (Liu et al., 1993), Tec1p (Bao et al., 2004; Bardwell, 2005; Chou et al., 2004; Madhani and Fink, 1997), Msa1p, and Msa2p (van der Felden et al., 2014), as well as loss of the repressive effects of negative regulators Dig1p and Dig2p (Bardwell et al., 1998; Cook et al., 1996; McCullagh et al., 2010). The net effect is transcriptional induction of target genes whose products bring about the filamentous cell type.

Cdc42p, Ste20p, Ste11p, and other proteins not only regulate the fMAPK pathway but also function in the HOG and mating pathways (Patterson et al., 2021; Saito and Posas, 2012; Saito and Tatebayashi, 2004; Vázquez-Ibarra et al., 2020; Wu et al., 2006). Each pathway has its own MAP kinase: Kss1p for the fMAPK pathway, Hog1p for HOG pathway, and Fus3p for the mating pathway (Hao et al., 2012). Kss1p and Fus3p are both phosphorylated during mating to modulate the mating response (Pryciak and Huntress, 1998). Each pathway also contains a pathway-specific scaffold. The well-studied scaffold Ste5p is an adaptor for the mating pathway (Malleshaiah et al., 2010). Ste5p binds to the MAPK Fus3p and catalytically unlocks the protein, making it available for phosphorylation by Ste7p (Good et al., 2009). Pbs2p [also the MAPKK (Posas and Saito, 1997)] and Ahk1p (Nishimura et al., 2016) are scaffolds for the HOG pathway, and Bem4p is a scaffold for the fMAPK pathway (Pitoniak et al., 2015). Other proteins also regulate the fMAPK pathway, such as the 14-3-3 proteins Bmh1p and Bmh2p (Roberts et al., 1997), although what role these proteins play in pathway specification remains incompletely explored.

We previously showed that Cdc42p can be degraded to modulate the activity of the fMAPK pathway (González and Cullen, 2022). By further examining potential regulatory mechanisms surrounding the Cdc42p-PAK module in this pathway, we found that Ste20p is also degraded by the 26S proteasome. Moreover, binding of Ste20p to Cdc42p stabilizes Ste20p protein levels. We propose that activation of the fMAPK pathway leads to turnover of Cdc42p and Ste20p to attenuate MAPK pathway signaling. We also identified a Ste20p-independent branch of the fMAPK pathway. The Ste20p-dependent branch required Bmh1p and Bmh2p, members of the 14-3-3 family of proteins, whereas the Ste20p-independent branch required the fMAPK pathway adaptor Bem4p. We also explored the role that other proteins that regulate the Cdc42p module play in MAPK pathway regulation. Some proteins primarily functioned in basal signaling, such as Rga1p, a GTPase-activating protein for Cdc42p that functions in the fMAPK pathway (Smith et al., 2002). Another protein, Rdi1p mainly impacted the pathway under activating conditions. We also found that the polarity adaptor Bem1p interacted with the tetra-span protein Sho1p, which provides a connection between a sensor that functions at the plasma membrane and proteins that regulate the GTPase module. These regulatory features deepen our understanding of MAPK pathway regulation in yeast and may extend to other Rho-dependent signaling pathways in eukaryotic organisms.

## RESULTS

### Ste20p is turned over in the 26S proteasome and is stabilized by interaction with Cdc42p

The fMAPK pathway is composed of proteins that function in an ordered manner, initiated by sensors at the plasma membrane, Msb2p and Sho1p (**Fig. 1A**). Msb2p and Sho1p function through Bem4p, Bem1p, and Rsr1p to activate the Cdc42p GTPase module. Similar to the mating (Lamson et al., 2002; Simon et al., 1995) and HOG (Raitt et al., 2000) pathways, the active or GTP-bound conformation of Cdc42p interacts with the PAK kinase Ste20p to promote activation of the fMAPK pathway kinase cascade (**Fig. 1A**). In the N-terminus of the protein, Ste20p contains a characteristic <u>C</u>dc42p- and <u>R</u>ac-interactive <u>b</u>inding motif (CRIB) that when bound to Cdc42p relieves auto-inhibition of the kinase domain of the protein (Ash et al., 2003; Lamson et al., 2002; Leberer et al., 1997).

**Figure 1.**
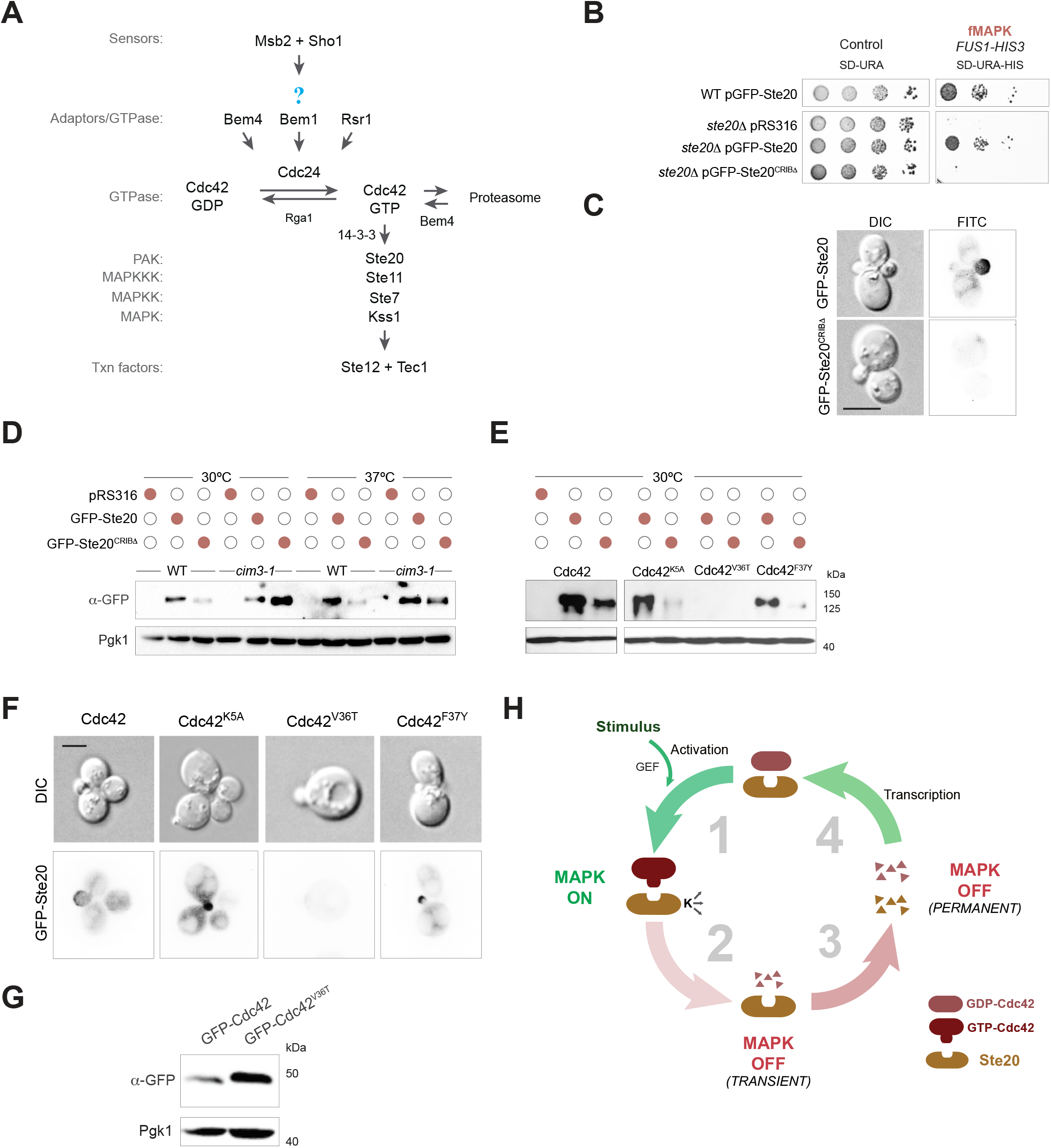
Role of the CRIB domain of Ste20p on the localization and turnover of the protein. **A)** The fMAPK pathway. Proteins known to regulate the fMAPK pathway are shown, question mark refers to how Sho1p connects to the GTPase module. **B)** Wild-type cells (WT, PC538, cells lack Ste4p) and cells lacking Ste20p (PC673) and expressing an empty plasmid (pRS316; PC2207), pGFP-Ste20p (PC4394), or pGFP-Ste20p^CRIB1′^ (PC4395) were spotted onto SD-URA and SD-URA-HIS media to evaluate the activity of the *FUS1-HIS3* growth reporter. Plates were incubated for 3 d at 30°C. **C)** Localization of GFP-Ste20p and GFP-Ste20p^CRIBΔ^ in wild-type cells and the *ste20Δ* mutant. Strains and plasmids in panel 1B were used. Scale bar, 5 µm. **D)** Immunoblot analysis of wild-type cells (PC5852) and the *cim3-1* mutant (PC5851) that contain pRS316 (CTL), pGFP-Ste20p (PC4394), and pGFP-Ste20p-CRIB (PC4395). Cells were grown at 30°C degree for 5 h and shifted to 37°C for 2h. **E)** Immunoblots were performed with wild-type cells (PC538) and cells harboring *CDC42* alleles K5A, V36T, and F37Y containing the same plasmids as in panel 1D. **F)** Strains described in panel 1E were analyzed by fluorescence microscopy. Scale bar, 5 µm. **G)** Level of GFP-Cdc42p (PC6454) and GFP-Cdc42p^V36T^ (PC7670) in wild-type cells measured by immunoblot analysis. **H)** Model showing the regulation between the GTPase Cdc42p and the PAK Ste20p. 1, under non-activating conditions, Cdc42p is in a GDP-bound conformation and does not interact with Ste20p. 2, environmental conditions (stimulus) induce GEF-dependent exchange of GDP-bound to GTP-bound Cdc42p which results in binding of Ste20p. Cdc42p-binding activates the kinase activity of Ste20p (K ->), resulting in rapid activation of the MAPK pathway (fast). 3, Ste20p may in some manner promote the degradation of GTP-Cdc42p, which results in a decrease in MAPK pathway activity. 4, Reduced levels of GTP-Cdc42p leads to free Ste20p, resulting in proteasomal degradation of Ste20p. Over time, transcription of the *CDC42* and *STE20* genes results in an increase in the levels of these proteins.

The activity of the fMAPK pathway can be evaluated by several tests, including by transcriptional reporters. A transcriptional reporter that normally functions in mating (*FUS1-HIS3*) provides a readout of fMAPK pathway activity in cells lacking an intact mating pathway [*ste4*, (Cullen et al., 2004)]. The wild-type strain (WT) and a strain lacking Ste20p (*ste20*Δ) were examined containing an empty plasmid (pRS316, URA3-marked) or plasmids containing genes encoding GFP-fusions to Ste20p (pGFP-Ste20p) or a version of Ste20p lacking the CRIB (334-369Δ) domain (pGFP-Ste20p^CRIBΔ^). Based on this reporter, cells lacking Ste20p are defective for induction of the *FUS1-HIS3* growth reporter and fail to grow on SD-URA-HIS media (**Fig. 1B**, *ste20*Δ pRS316). As expected, introduction of pGFP-Ste20p restored MAPK pathway activity and growth to the *ste20*Δ mutant. However, pGFP-Ste20p^CRIBΔ^ was defective for fMAPK pathway activity (**Fig. 1B**). These results show that in line with published results, Ste20p interact with Cdc42p by its CRIB domain to induce the fMAPK pathway.

Cdc42p binding to Ste20p is also required for Ste20p localization in buds and presumptive bud sites (Ash et al., 2003; Lamson et al., 2002; Moran et al., 2019). Accordingly, in wild type cells, GFP-Ste20p was found in buds by fluorescence microscopy (**Fig. 1C**). By comparison, GFP-Ste20p^CRIBΔ^ showed a diffuse pattern (**Fig. 1C**). To further explore this phenotype, immunoblot analysis was performed using antibodies to the GFP epitope. Immunoblot analysis showed that GFP-Ste20p^CRIBΔ^ was present at lower levels than GFP-Ste20p (**Fig. 1D**, WT, 30°C). This unexpected finding indicates that the CRIB domain is required to maintain normal levels of Ste20p in the cell.

Protein levels can be impacted by turnover, such as by the ubiquitination and degradation of proteins by the 26S proteasome (Grice and Nathan, 2016). In line with this possibility, the level of GFP-Ste20p^CRIBΔ^ was higher in the *cim3-1* mutant (**Fig. 1D**, *cim3-1*, 30°C), which contains a conditional mutation in the gene encoding the proteasomal ATPase Rpt6p, a protein required for 26S proteasome function (Ghislain et al., 1993). Elevated levels of GFP-Ste20p^CRIBΔ^ were observed in the *cim3-1* mutant at 30°C and 37°C (**Fig. 1D**, 37°C), which indicates that a partial defect in proteasome function is sufficient to restore normal levels of Ste20p in the cell. By comparison, the level of wild-type GFP-Ste20p was not impacted in the *cim3-1* mutant at 30°C or 37°C. These results reveal a new aspect of Ste20p regulation, by turnover of a version of the protein that is defective for interaction with Cdc42p in the 26S proteasome.

Ste20p lacking its CRIB domain might be unstable and degraded because the protein is misfolded. Alternatively, Ste20p might be protected from turnover by binding to Cdc42p. To explore these possibilities, cells expressing versions of Cdc42p compromised for specific functions as the sole copy in the cell were examined. In particular, Cdc42p^V36T^ and Cdc42p^F37Y^ contain amino acid substitutions in the switch I domain, which is required for binding proteins with CRIB motifs, including Ste20p [other effectors include Cla4p, Gic1p, and Gic2p (Kozminski et al., 2000; Moskow et al., 2000)]. As controls, wild-type Cdc42p and a version of the protein that contains an amino acid change in a domain unrelated to its interaction with effector proteins (Cdc42p^K5A^) were also examined. Compared to controls (Cdc42 and Cdc42^K5A^), cells expressing Cdc42p^V36T^ and to a lesser degree Cdc42p^F37Y^ showed reduced levels of the wild-type version of GFP-Ste20p by immunoblot analysis (**Fig. 1E**, Cdc42p^V36T^). By fluorescence microscopy, the level of GFP-Ste20p was also reduced in cells expressing Cdc42p^V36T^ (**Fig. 1F**, Cdc42p^V36T^). Cdc42p^F37Y^ also showed reduced levels of GFP-Ste20p but not to the same degree Cdc42p^V36T^ (**Fig. 1E**) and showed a normal localization pattern (**Fig. 1F**). This result is consistent with the fact that Cdc42p^V36T^ exhibits a more severe defect in fMAPK pathway activity compared to Cdc42p^F37Y^ (Basu et al., 2020). These results are also consistent with the fact that Cdc42p^V36T^ exhibits a more striking defect for interaction with Ste20p than Cdc42p^F37Y^ (Gladfelter et al., 2001). These results independently validate the idea that the interaction between Cdc42p and Ste20p is important for maintaining normal levels of Ste20p in the cell.

We previously showed that the active or GTP-bound conformation of Cdc42p is preferentially turned over in the proteosome compared to the wild-type conformation, which is inhibited by the adaptor Bem4p [**Fig. 1A**, (González and Cullen, 2022)]. One hypothesis to explain this turnover mechanism is that the active conformation of Cdc42p interacts with effector proteins that function in some manner to promote turnover of the G-protein. To test this possibility, the level of the Cdc42p^V36T^ protein was also examined. GFP-Cdc42p^V36T^ was present in the cell at higher levels than wild-type GFP-Cdc42p (**Fig. 1G**). The level of Cdc42p was also higher in cells lacking Ste20p (González and Cullen, 2022). This result could be interpreted to mean that the interaction of Cdc42p with effector proteins like Ste20p results in turnover of the protein.

Together with the above results, these observations suggest a model for turnover of the Rho-PAK module (**Fig. 1H**). In step 1, a stimulus induces GEF-dependent activation of Cdc42p. The active or GTP-bound conformation of Cdc42p binds and activates effector proteins including Ste20p. In step 2, GTP-Cdc42p is turned over by degradation in the 26S proteosome, which may be stimulated by binding to Ste20p or other effector proteins. In step 3, Ste20p molecules that no longer bind to Cdc42p are themselves degraded. The disappearance of GTP-bound Cdc42p and Ste20p may result in a more permanent shut off of the fMAPK pathway (and mating and HOG pathways, although we have not tested these possibilities). In step 4, the transcription of genes encoding *CDC42* and *STE20* return the proteins to their baseline states. Thus, a balance of stability/turnover of Rho-PAK adds a new layer of regulation to MAPK pathway signaling.

### A Cdc42p-dependent branch of the fMAPK pathway functions outside of Ste20p and requires the adaptor protein Bem4p

We previously identified a turnover-defective (TD) and GTP-locked (Q61L) version of Cdc42p that hyperactivates the fMAPK pathway [Cdc42p^Q61l+TD^ (González and Cullen, 2022)]. This tool allowed evaluation of the fMAPK pathway activity in different genetic contexts. As has been reported (González and Cullen, 2022), Cdc42p^Q61L+TD^ induced the fMAPK pathway to slightly higher levels than GTP-locked Cdc42p, based on the *FUS1-HIS3* growth reporter (**Fig. 2A**). Surprisingly, Cdc42p^Q61L+TD^, and to some degree Cdc42p^Q61L^, induced the fMAPK pathway in cells lacking Ste20p (**Fig. 2A**, *ste20*Δ). Because this observation was unexpected, the activity of the fMAPK pathway was assessed by additional reporters of the fMAPK pathway. By another fMAPK pathway-dependent transcriptional reporter, *FRE-lacZ* (Madhani et al., 1999), Cdc42p^Q61L+TD^ also partially bypassed the requirement for Ste20p (**Fig. 2B**,*ste20*Δ). Cdc42p^Q61L^ appears to strongly hyperactivate the *FRE-lacZ* reporter compared to *FUS1-HIS3* (Q61L compare **Fig. 2A** and **Fig. 2B**); however, use of the competitive inhibitor 3-amino-1,2,4 triazole (ATA) showed that the reporters behaved similarly (*Fig. S1A*). Cdc42p^Q61L+TD^ did not bypass the requirement for the MAPKKK Ste11p (**Fig. 2B**, *Fig. S1A, ste11*Δ), indicating that Cdc42p signals independently of Ste20p and dependent on Ste11p. The Ste11p-interacting protein Ste50p (Ramezani Rad et al., 1998; Sharmeen et al., 2019; Xu et al., 1996) and transcription factor Ste12p were also required for signaling induced by Cdc42p^Q61L+TD^ (*Fig. S1A*). The activity of the fMAPK pathway was also measured by phosphorylation of the MAP kinase, Kss1p, which showed the same results (**Fig. 2C**, see quantitation below the graph). We note that the level of P∼Kss1p in cells lacking Ste20p was reduced compared to wild-type cells (**Fig. 2C**, WT Cdc42^Q61L+TD^), which as seen for *FRE-lacZ,* is indicative of a partial bypass. Activation of the fMAPK pathway induces the formation of elongated cells that are capable of invasive/pseudohyphal growth. Cdc42p^Q61L+TD^ induced cell elongation in cells lacking Ste20p, which was dependent on Ste11p (**Fig. 2D**). The fMAPK pathway also induces invasive growth that can be examined by the plate-washing assay (Roberts and Fink, 1994). Cells lacking Ste20p showed detectable invasive growth when expressing Cdc42p^Q61L^ and Cdc42p^Q61L+TD^ (*Fig. S1B*). These observations together indicate that Cdc42p regulates a Ste20p-independent branch of the fMAPK pathway.

**Figure 2.**
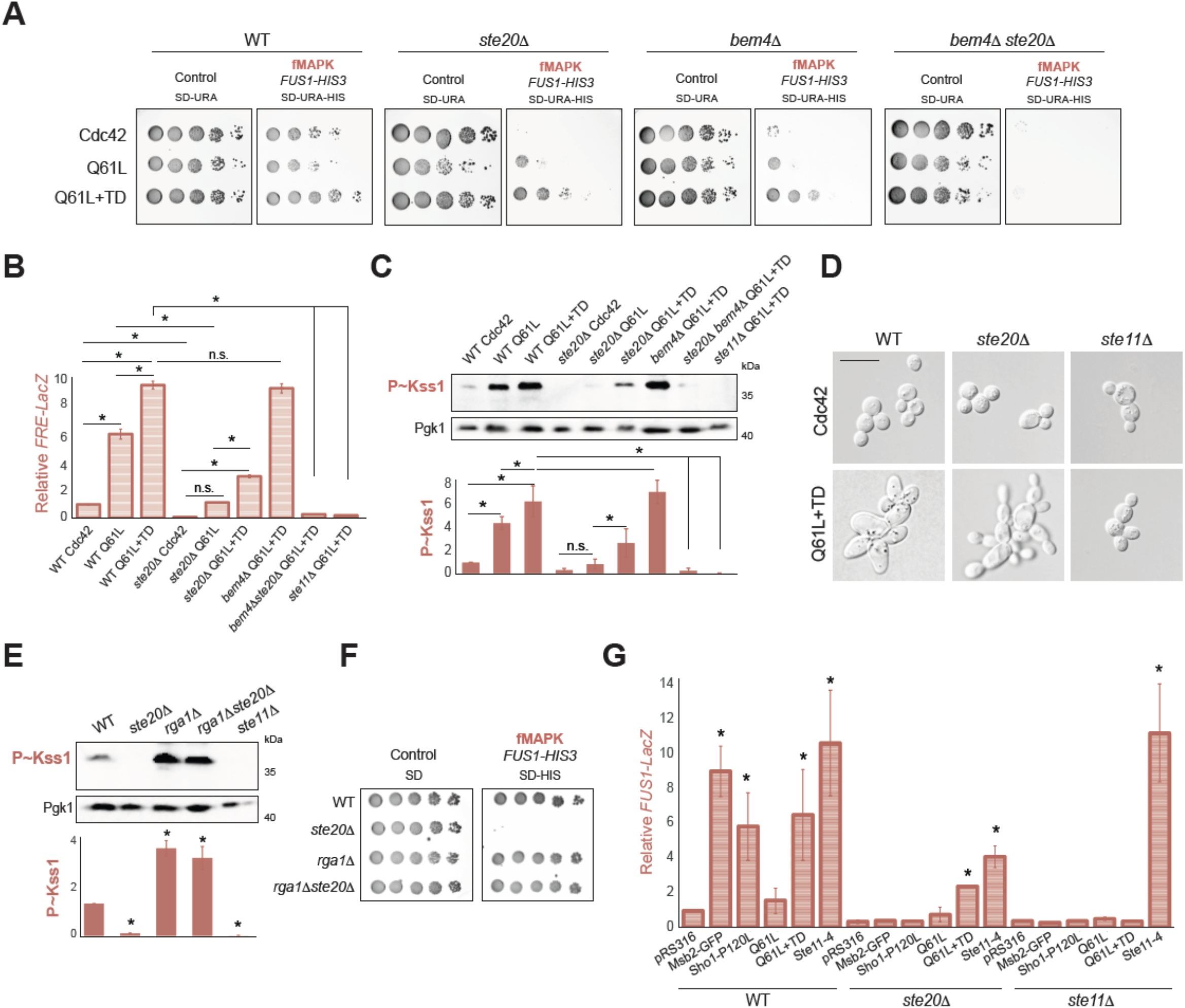
fMAPK pathway activity independent of Ste20p. **A)** Wild-type cells and the indicated mutants expressing pGFP-Cdc42p (Cdc42; PC6454), pGFP-Cdc42p^Q61L^ (Q61L; PC7458) or pGFP-Cdc42p^Q61L+TD^ (Q61L+TD: PC7654) were grown on SD-URA or SD-URA-HIS media. See Fig. 1B for details. **B)** Δ-galactosidase assay of the *FRE-LacZ* transcriptional reporter to analyze fMAPK pathway activity. Wild-type cells (WT; 6810), the *ste20*Δ (PC7772) mutant expressing same plasmids explained in panel 2A. Cells in panel B were grown for 5 h in SD-URA-LEU media. Values show the relative transcriptional activation compared to wild-type cells expressing Cdc42. Error bars represent the SD from three biological replicates (n =3). Data were analyzed by one-way ANOVA, and P values (asterisk < 0.05) were obtained using Tukey’s multiple comparison test. **C)** Levels of P∼Kss1p of same strains as described in Fig. 2B grown for 6 h in SD-URA media. Error bars represent the SD from three biological replicates (n= 3). Data were analyzed by one-way ANOVA, and P values (asterisk < 0.05) were obtained using Tukey’s multiple comparison test. **D)** Microscopic examination of wild-type cells, and the s*te20*Δ and *ste11*Δ mutans expressing wild-type Cdc42p (Cdc42) and Cdc42p^Q61L+TD^ (Q61L+TD) grown for 24 h on SD-URA media. Scale bar, 15 µm. **E)** Levels of P∼Kss1p of wild-type strains and the indicated mutants grown for 5 h in SD-URA media. Error bars represent SD from three biological replicates (n=3). Data were analyzed by one-way ANOVA, and P values (asterisk < 0.05) were obtained using Tukey’s multiple comparison test. **F)** Wild-type cells and the indicated mutants were grown on SD or SD-HIS media to study the activity of *FUS1-HIS3* growth reporter. See Fig. 1B for details. **G)** Wild-type cells, and the s*te20*Δ and *ste11*Δ mutans expressing pRS316 (PC2207), Msb2-GFP (PC1696), Sho1pP120L (PC1715), Cdc42p^Q61L^, Cdc42p^Q61L+TD^, or Ste11-4 (PC1441) were grown in SGAL-URA media to evaluate the activity of the *FUS1-LacZ* reporter. Error bars represent SD from two biological replicates (n= 2). Data were analyzed by one-way ANOVA, and P values (asterisk < 0.05) were obtained using Tukey’s multiple comparison test.

As a separate test, Cdc42p-dependent signaling was examined in cells lacking Rga1p, which is one of the GTPase activating protein (GAPs) for Cdc42p, and the main GAP that functions in the fMAPK pathway (Smith et al., 2002). Cells lacking Rga1p show higher levels of GTP-Cdc42p (Tong et al., 2007). Consistent with this idea, the *rga1*Δ mutant induced higher levels of P∼Kss1p than wild-type cells (**Fig. 2E**, *rga1*Δ). Cells lacking Rga1p and Ste20p showed similar levels of P∼Kss1p than cells lacking Rga1p alone (**Fig. 2E**, *rga1*Δ *ste20*Δ). The *rga1*Δ *ste20*Δ double mutant also rescued the signaling defect of the *ste20*Δ mutant based on the *FUS1-HIS3* reporter (**Fig. 2F**). Altogether, these two different types of experiments support the idea that Cdc42p also regulates a Ste20p-independent branch of the fMAPK pathway.

The adaptor protein Bem4p interacts with Cdc42p and Ste11p (Basu et al., 2020; Pitoniak et al., 2015) and might mediate signaling in the Ste20p-independent branch. Consistent with this possibility, Cdc42p^Q61L+TD^ did not bypass the signaling defect of the *ste20*Δ *bem4*Δ double mutant, by *FUS1-LacZ* (**Fig. 2A**), *FRE-LacZ* (**Fig. 2B**), or P∼Kss1p analysis (**Fig. 2C**). As previously reported (González and Cullen, 2022), Cdc42p^Q61L+TD^ bypassed the signaling defect of the *bem4*Δ single mutant (**Fig. 2**, **A**, **B** and **C**). Therefore, Cdc42p induces Ste20p-independent signaling through Bem4p.

Ste20p is a common component that is shared between three Cdc42p-dependent MAPK pathways (fMAPK, mating, and HOG), so we tested whether Cdc42p^Q61L+TD^ bypassed the requirement for Ste20p in the HOG and mating pathways. To assess the activity of the HOG pathway, cells lacking the Sln1p branch of the HOG pathway (*ssk1*Δ) were examined, which is redundant with the Ste11p branch (Saito and Posas, 2012). Cells expressing Cdc42p^Q61L+TD^ that also lacked Ste20p did not grow in high-osmolarity media (*Fig. S1C*), which provides a readout of the HOG pathway (Maeda et al., 1995). Similarly, cells expressing Cdc42p^Q61L+TD^ did not respond to α-factor in cells lacking Ste20p (*Fig. S1D*), which provides a readout of the mating pathway (Sprague et al., 1983). Therefore, Cdc42p^Q61L+TD^ may preferentially stimulate the Ste20p-independent branch of the fMAPK pathway.

Hyperactive versions of sensors for the fMAPK pathway, Msb2p [GFP-Msb2, (Adhikari et al., 2015)] and Sho1p [Sho1p^P120L^, (Vadaie et al., 2008)] have also been useful tools to examine signal flow through the fMAPK pathway. Hyperactive versions of these proteins did not bypass the requirement for Ste20p in the fMAPK pathway, based on *FUS1-lacZ* (**Fig. 2G**) and *FUS1-HIS3* analysis (*Fig. S1E*). A plasmid expressing the hyperactive allele *STE11-4* was used as a control. Msb2p and Sho1p may selectively regulate the Ste20p branch of the fMAPK pathway. To summarize, the fMAPK pathway is composed of Ste20p-dependent and Ste20p-independent branches that are regulated by different combinations of proteins.

### 14-3-3 proteins Bmh1p and Bmh2p regulate the Ste20p branch of the fMAPK pathway

Bmh1p and Bmh2p are members of the 14-3-3 family of proteins (van Hemert et al., 2001) which were previously identified as regulators of the fMAPK pathway that interact with and function through Ste20p (Roberts et al., 1997). Bmh1p and Bmh2p were identified by our laboratory in a genome-wide screen for proteins that when overexpressed stimulate the activity of the fMAPK pathway (Jamalzadeh et al., 2020). Although these proteins were found to associate with Ste20p, their role in regulating the three Ste20p-dependent pathways has yet to be fully explored (**Fig. 3A**). To confirm a role for Bmh1p and Bmh2p proteins in regulating the fMAPK pathway, cells containing plasmids overexpressing the *BMH1* or *BMH2* genes were examined by the *FUS1-HIS3* growth reporter. Based on this test, overexpression of the *BMH* genes stimulated the fMAPK pathway (**Fig. 3B**). Overexpression of these genes also induced a growth defect, which did not result from elevated fMAPK pathway activity as the growth defect was also seen in cells lacking an intact fMAPK pathway (**Fig. 3B**, *ste11*Δ). Similarly, deletion of the *BMH1* or *BMH2* genes caused a decrease in *FUS1-HIS3* reporter activity (*Fig. S2A*). We were unsuccessful in disrupting both genes, which is lethal in some backgrounds (van Heusden et al., 1995; van Heusden and Steensma, 2006), and which has previously been shown to cause a defect in filamentous growth (Roberts et al., 1997). Overexpression of *BMH1* or *BMH2* also induced invasive growth (**Fig. 3C**, Washed), and overexpression of *BMH1* induced an elongated cell morphology by microscopy (**Fig. 3D**, Bmh1p and Bmh2p behaved similarly). Genetic suppression (or epistasis) analysis showed that overexpression of *BMH1* and *BMH2* induced the *FUS1-HIS3* reporter in a manner that was dependent on Ste20p and Ste11p (**Fig. 3B**). Similarly, the filamentous cell morphology induced by *BMH1* was dependent on Ste20p and Ste11p (**Fig. 3D**). These results indicate that the Bmh proteins function in the Ste20p branch of the fMAPK pathway.

**Figure 3.**
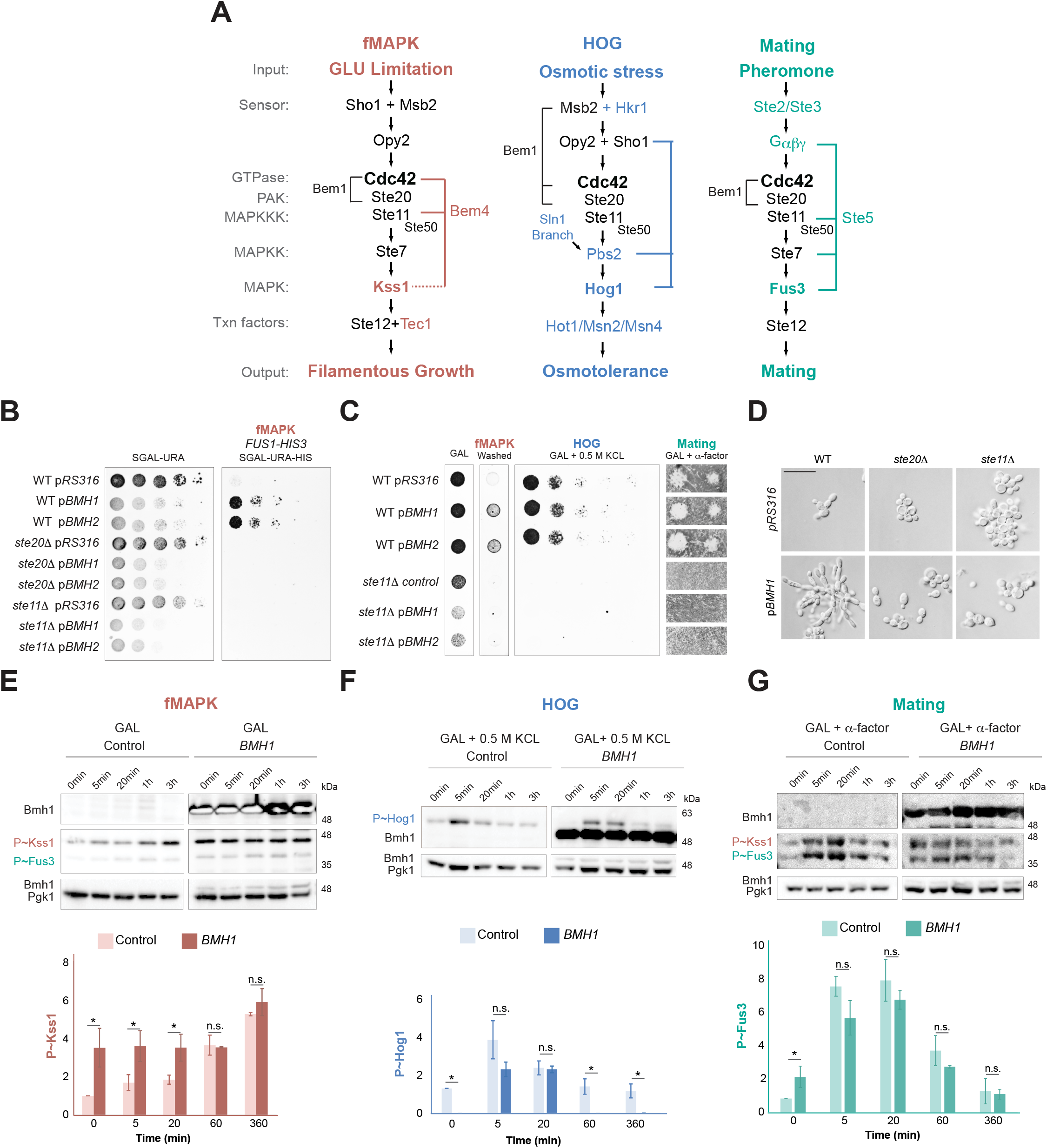
The impact of Bmh1p and Bmh2p on fMAPK, HOG, and mating pathway activity. **A)** Proteins that regulate the fMAPK (red), HOG (blue) and mating (green) pathways, common regulators are colored in black. **B)** Wild-type cells (PC538), and the *ste20*Δ (PC673) and *ste11*Δ (PC3862) mutants expressing pRS316, pP*_GAL1_*-BMH1 or pP*_GAL1_*-BMH2 (Gelperin et al., 2005) were grown on SGAL media to study the activity of the *FUS1-HIS3* reporter. See Fig. 1B for details. **C)** Wild-type cells (PC6810), and the *ste20*Δ (PC7772) and *ste11*Δ (PC6604) in the *ssk1*Δ background expressing same plasmids as described in panel 3A were grown on YEPGAL (Gal), YEPGAL supplemented with 0.5 M KCl or YEPGAL supplemented with 6 µM or 1.6 µM of α-factor. **D)** Microscopic examination of same cells described in panel 3A grown for 6 h in SGAL media. Scale bar, 30 µm. **E)** The *ssk1*Δ mutant expressing pRs316 (control) or pP*_GAL1_*-BMH1 was grown during the indicated time points in YEPGAL media. Immunoblots were probed with p44/42 antibodies to detect P∼Kss1p and P∼Fus3p and with α-Pgk1p antibodies. Error bars represents SD from two biological replicates (n= 2). Data were analyzed by one-way ANOVA, and P values (asterisk < 0.05) were obtained using Tukey’s multiple comparison test. **F)** The *ssk1*Δ mutant expressing pRS316 (control) or pP*_GAL1_*-BMH1 was grown during the indicated time points in YEPGAL media supplemented with 0.5 m KCl. Immunoblots were probed with p38 antibodies to detect P∼Hog1p and with α-Pgk1 antibodies. Data were analyzed as described in panel 3E. **G)** The *ssk1*Δ mutant expressing pRS316 (control) or pP*_GAL1_*-BMH1 was grown during the indicated time points in YEPGAL media supplemented with 1.6 µM of α-factor. See Fig. 3E for details. Data were analyzed as described in panel 3E.

The Bmh1p and Bmh2p proteins might specifically regulate the fMAPK pathway or function as general regulators of Cdc42p-dependent MAPK pathways that share components (**Fig. 3A**). Overexpression of the *BMH1* or *BMH2* genes did not influence growth in media supplemented with salt (**Fig. 3C**, 0.5 M KCl) and did not impact mating, based on sensitivity to pheromone (**Fig. 3C**, GAL + α-factor). These results indicate that the Bmh1p and Bmh2p proteins do not play a major role in regulating the HOG or mating pathways. The phosphorylation of the MAP kinases for each of the Cdc42p-dependent MAP kinase pathways was also examined. To examine basal and activated conditions, cells were grown in media containing galactose over a time-course (**Fig. 3E**, control, 1h; 3h), which induces phosphorylation of the fMAPK Kss1p (Basu et al., 2020; González and Cullen, 2022; Prabhakar et al., 2021). As reported (Basu et al., 2020; Jamalzadeh et al., 2020) P∼Kss1p levels increased in galactose over time. Overexpression of *BMH1* induced high levels of P∼Kss1p over the entire time course of the experiment (**Fig. 3E**, *BMH1*, 0 min through 3h). This result indicates that overproduction of Bmh1p induces the fMAPK pathway. Under inducing conditions (3hr), *BMH1* did not cause a further increase in P∼Kss1 levels above its maximal levels (**Fig. 3E**, graph, 3h), which may indicate that Bmh1p preferentially impacts the basal-to-activated signaling of the fMAPK pathway.

A similar approach was used to evaluate the role of Bmh1p in regulating the HOG pathway. Phosphorylation of the MAPK Hog1p was assessed by exposure of cells to salt over time, which results in the rapid increase in P∼Hog1p levels (**Fig. 3F**). Overexpression of *BMH1* did not induce phosphorylation P∼Hog1p at any time point tested (**Fig. 3F**, *BMH1*, 0 min). In fact, the level of P∼Hog1p was lower for several time points especially in no-salt environments (t=0) where basal levels of P∼Hog1p were seen to be reduced. Therefore, overexpression of Bmh1p might result in a slight reduction in HOG pathway activity. In response to the mating pheromone α-factor, overexpression of *BMH1* did not induce the phosphorylation of Fus3p (**Fig. 3G**). The level of P∼Fus3p was in fact reduced compared to control conditions, indicating that Bmh1p overexpression may slightly attenuate the mating pathway. This may be a modest effect, as cells overexpressing *BMH1* were able to produce shmoos (*Fig. S2B*). Taken together, these results show that Bmh1p and Bmh2p regulate the Ste20p branch of the fMAPK pathway. Because overexpression of the Bmh1p and Bmh2p proteins did not have a major impact on the HOG or mating pathways, these proteins might not play a major role in mediating specificity between Cdc42p-dependent pathways.

### Cdc24p phosphorylation and positive feedback are not major determinants of fMAPK pathway regulation

Rho GTPases are activated by guanine nucleotide exchange factors (GEFs) that promote exchange from GDP to GTP and are inactivated by GTPase activity by <u>G</u>TPase <u>a</u>ctivating <u>p</u>roteins (GAPs). In addition, Rho GTPases are regulated by RhoGDIs that can inhibit Rho proteins by extracting them from membranes. In this and the following two sections, the role that GEF modifications, GAPs, and Rho GDIs play in regulating fMAPK pathways were examined (**Fig. 4A**).

**Figure. 4.**
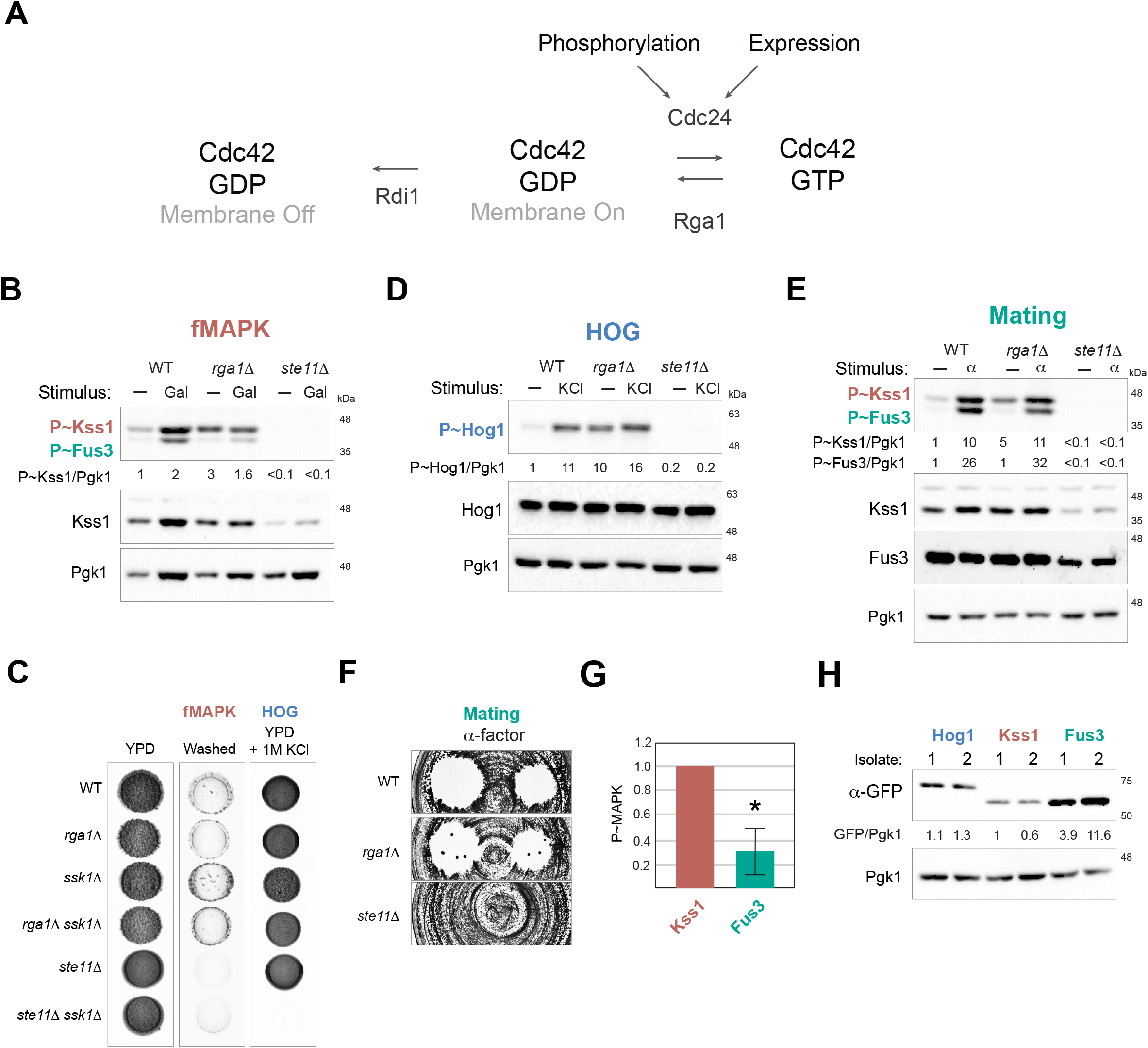
Rga1p regulates basal activity of Cdc42-dependent MAPK pathways. **A)** The regulation of Cdc42p by Cdc24p, Rga1p, and Rdi1p. **B)** Wild-type cells (PC6810) and the *rga1*Δ (PC6687) and *ste11*Δ (PC6604) mutants form the *ssk1*Δ background were grown in YPD (-) or YPGAL (Gal) for 5.5 h at 30°C. Immunoblots were probed with p44/42 antibodies to detect P∼Kss1p and P∼Fus3p, α-Kss1p and α-Pgk1p antibodies. P∼Kss1/Pgk1 ratio refers to relative levels of P∼Kss1p to Pgk1p of the blot shown. **C)** Wild-type cells and mutants indicated were grown in YPD media for 3 d at 30°C (YPD) and the colony grown was washed (fMAPK). Same cells were grown in YPD supplemented with 1M KCl for 3 d at 30°C (HOG). **D)** Same cells described in panel 4A were grown in YPD (-) or YPD supplemented with 1M KCL (KCl) for 5 min at 30°C. Immunoblots were probed with p38 antibodies to detect P∼Hog1p, α-Hog1p and α-Pgk1p antibodies. P∼Hog1/Pgk1 ratio refers to relative levels of P∼Hog1p to Pgk1p of the blot shown. **E)** Same cells described in panel 4A were grown in YPD (-) or YPD supplemented with 6 µM of α-factor (α) for 5 min at 30°C. Immunoblots were probed with p44/42 antibodies to detect P∼Kss1p and P∼Fus3p, α-Kss1p, α-Fus3p and α-Pgk1p antibodies. P∼Kss1/Pgk1 ratio refers to relative levels of P∼Kss1p to Pgk1p of the blot shown. P∼Fus3/Pgk1 ratio refers to relative levels of P∼Fus3p to Pgk1p of the blot shown. **F)** Halo formation in response to α-factor of same cells described in panel 4B. Cells were spread on YPD media and α-factor was spotted at two concentrations on the surface, 6 and 2 μM, to study cell-cycle arrest. **G)** Average levels of the indicated P∼MAPKs based on multiple independent biological replicates. Band intensity was quantitated by ImageJ. P∼Kss1p levels were set to a value of 1, and average values of P∼Fus3p are reported. Error bars represent the standard difference between >12 experiments. P-value < 0.05. **H)** Levels of GFP-tagged MAP kinases. Immunoblots were performed as in panel 4B, except anti-GFP antibodies were used.

We first examined Cdc24p phosphorylation. Cdc24p is phosphorylated in a cell-cycle dependent manner (Bose et al., 2001; Gulli et al., 2000; Rossman et al., 2005). Cdc24p is required for fMAPK pathway activity (Pitoniak et al., 2015; Prabhakar et al., 2020), and cells expressing a temperature-sensitive allele, *cdc24-4*, are not able to induce the fMAPK pathway, even at the permissive temperature (*Fig. S3A*). Mass spectrometry analysis identified residues on the Cdc24p protein that are phosphorylated in several domains of the protein (Wai et al., 2009). Versions of Cdc24p containing substitutions of known phosphorylation sites, expressed from *LEU2*-based plasmids (Wai et al., 2009) were tested in cells in which the genomic copy of *CDC24* was disrupted with the antibiotic resistance marker *NAT* (*cdc24::NAT*) that also contained the wild-type *pCDC24 URA3* plasmid. The wild-type *pCDC24 URA3* plasmid was forced out of cells by growth on 5-fluoroorotic acid (5-FOA) to evaluate phosphosite mutants of Cdc24p. Versions tested included serine and threonine substitutions in the PH domain (PH), substitutions in a linker domain (Linker) connecting the PH and PB1 domains, substitutions in both domains (PH/Linker), and a version with substitutions in all 35 phosphorylation sites (T35A). All versions tested rescued the viability defect of the *cdc24::NAT* mutant, based on growth on 5-FOA, and retained function in the fMAPK pathway based on the *FUS1-HIS3* growth reporter (*Fig. S3B*) and phospho-immunoblot analysis (*Fig. S3C*). The results indicate that phosphorylation of Cdc24p is not critical for regulation of the fMAPK pathway.

In *Candida albicans*, expression of the *CDC24* gene is induced during hyphal growth as a transcriptional target of the analogous Cek MAPK pathway (Bassilana et al., 2003). A transcriptional reporter for *CDC24* (p*CDC24-lacZ*) was constructed and tested for dependency on the fMAPK pathway. Based on this reporter, *CDC24* expression was not dependent on the fMAPK pathway (*Fig. S3D*). This result is consistent with expression profiling data from our laboratory (Adhikari and Cullen, 2014; Chavel et al., 2014; Chow et al., 2019) and with the fact that overexpression of *CDC24* does not induce fMAPK pathway activity or invasive growth in *S. cerevisiae* (Jamalzadeh et al., 2020). Thus, phosphorylation of Cdc24p and positive feedback are not major determinants in the regulation of the fMAPK pathway.

### Rga1 regulates basal but not active levels of Cdc42p-dependent MAPK pathways

We next examined the role of Rga1p. As mentioned above, Rga1p is the main GAP for Cdc42p in the fMAPK pathway (Smith et al., 2002) (**Fig. 4A**), and as shown above, cells lacking Rga1p showed elevated levels of P∼Kss1p (**Fig. 2E**). To more thoroughly examine the role of Rga1p, MAPK pathway was examined under basal (glucose replete) and pathway activated (galactose) conditions. Under glucose-replete conditions, the *rga1*Δ mutant showed elevated levels of P∼Kss1p (**Fig. 4B**, minus sign refers to glucose-replete conditions, compare WT to *rga1*Δ). These results show that Rga1p attenuates the fMAPK pathway under basal or non-activating conditions. By comparison, Rga1p did not dampen fMAPK pathway activity under activating conditions (**Fig. 4B**, Gal, compare WT to *rga1*Δ). This result was unexpected because Rga1p is expected to function under conditions when Cdc42p is active in the fMAPK pathway. These results were supported be phenotypic examination, which showed that cells lacking Rga1p showed a similar pattern of invasive growth as wild type (**Fig. 4C**). Therefore, Rga1p does not play a major role in regulating the fMAPK pathway under conditions when the pathway is active and cells are undergoing invasive growth.

Rga1p has previously been shown to inhibit the pheromone response pathway and dampen crosstalk between the HOG and mating pathways (O’Rourke and Herskowitz, 1998; Stevenson et al., 1995). Like for the fMAPK pathway, Rga1p also dampened basal HOG pathway activity (**Fig. 4D**, no stimulus designated by minus sign, compare WT to the *rga1*Δ mutant). However, Rga1p had a modest effect on the activated HOG pathway when cells were induced by osmotic stress (**Fig. 4D**, KCl compare WT to *rga1*Δ). Similarly, Rga1p did not impact osmotolerance, as seen in cells lacking the redundant Sln1p branch of the HOG pathway (**Fig. 4C**, *ssk1*Δ). Cells lacking Rga1p also showed higher basal mating pathway activity (**Fig. 4E**, no stimulus as designated by minus sign, compare WT to *rga1*Δ) but did not impact P∼Fus3p levels after treatment with α-factor (**Fig. 4E**). Similarly, the growth arrest in response to α-factor treatment, which also requires the mating pathway, was not impacted in cells lacking Rga1p (**Fig. 4F**). Rga1p therefore negatively regulates Cdc42p-dependent MAPK pathways mainly under basal conditions. Under activating conditions, Rga1p does not play a major role.

In wild-type cells and lacking Rga1p, we noticed that the level of P∼Kss1p was higher than the level of P∼Fus3p [e.g. (**Figs. 4B** and **4E**)]. This observation does not result from one kinase being preferentially recognized over another by the phosphor-antibodies, because addition of α-factor stimulated phosphorylation of both kinases equally (**Figs. 3G** and **4E**) as reported (Basu et al., 2020; Prabhakar et al., 2021). To confirm this result, the ratio of P∼Kss1p/P∼Fus3p was quantitated over a panel of phospho-immunoblots (**Fig. 4G**). Higher levels of basal P∼Kss1p than P∼Fus3p could result from relatively higher levels of the Kss1p protein in cells. GFP fusions to MAP kinase proteins showed however that Kss1p-GFP was present at lower levels than Fus3p-GFP or Hog1p-GFP (**Fig. 4H**). P∼Kss1p levels may be higher than P∼Fus3p levels because phosphorylation of Fus3p is dependent upon binding by Ste5p, which catalytically unlocks the protein (Good et al., 2009). These results demonstrate that the basal activity of Cdc42p-dependent MAPK pathways differ from each other and is not determined solely by the level of MAP kinase protein in the cell.

### Role of Rdi1p in regulating MAPK pathway signaling

Rho GTPases are also regulated by Rho GDP-dissociation inhibitors (GDIs) which control their subcellular localization (Boulter et al., 2010; Golding et al., 2019). The sole guanine nucleotide dissociation inhibitor (GDI) in budding yeast, Rdi1p, extracts Cdc42p from the plasma membrane and retains it in cytosolic pools preventing its activation by Cdc24p (Johnson, 1999; Richman et al., 2004) (**Fig. 4A**). As cells lacking *RDI1* had subtle phenotypes, the activity of MAPK pathways was examined in cells overexpressing *RDI1* by a galactose-inducible promoter. Overexpression of *RDI1* would be expected to promote Cdc42p extraction from membranes and thereby cause a decrease in MAPK pathway activity. As expected, overexpression of *RDI1* caused a reduction in fMAPK pathway activity under basal (minus, glucose) and activating conditions (Gal) (**Fig. 5A**). Functional tests showed that *RDI1* overexpression also caused a reduction in invasive growth (**Fig. 5B**, washed). With respect to other MAPK pathways, overexpression of *RDI1* also dampened the HOG (**Fig. 5C,** 1.4-fold) and Mating (**Fig. 5D,** 1.5-fold) pathways but to a lesser degree than fMAPK (**Fig. 5A**, 3.2-fold). Consistently, the fMAPK pathway was more affected that HOG and mating pathways by functional tests (**Fig. 5, B** and **E**). The interaction between Bem4p and Cdc42p is required for fMAPK pathway activation (Basu et al., 2020). Overexpression of *RDI1* by a galactose-inducible promoter reduced the interaction between Cdc42p and Bem4p by two-hybrid analysis (**Fig. 5F**). These results indicate that Cdc42p levels at the plasma membrane are critical for MAPK pathway signaling. The requirement for Cdc42p levels at the plasma membrane are different for different MAPK pathways. Moreover, the results show that the fMAPK pathway is highly sensitive to Cdc42p at the plasma membrane for activation.

**Figure 5.**
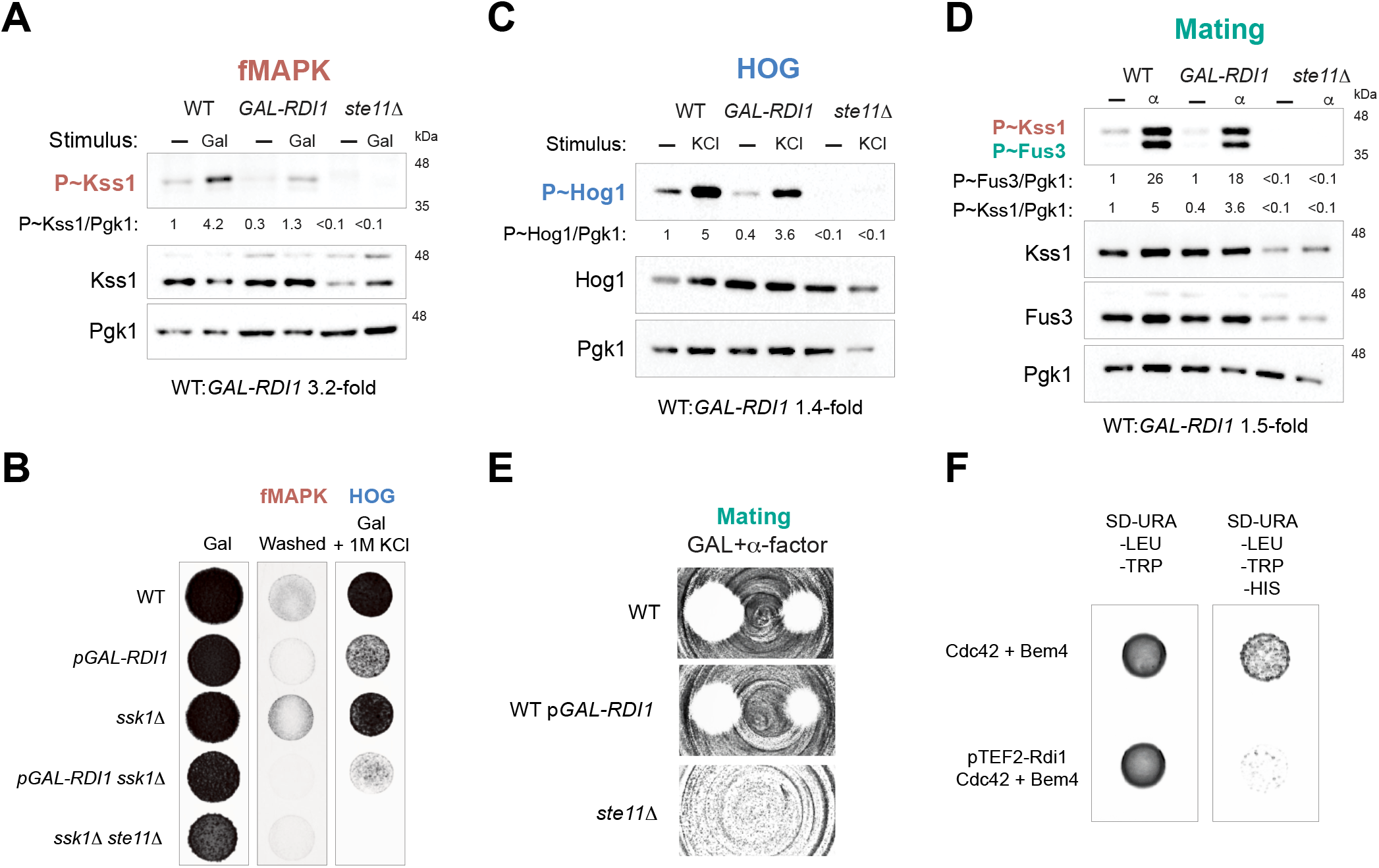
*RDI1* overexpression inhibits the activity of Cdc42-dependent MAPK pathways. **A)** Wild-type cells, or cells overexpressing *RDI1*, and the *ste11*Δ mutant (all in the *ssk1*Δ background) were grown in YPD (-) or YP-GAL (Gal) media for 5.5 h at 30°C. Immunoblots were probed with p44/42 antibodies to detect P∼Kss1p and P∼Fus3p, as well as α-Kss1p and α-Pgk1p antibodies as indicated. The P∼Kss1/Pgk1 ratio refers to relative levels of P∼Kss1p to Pgk1p of the blot shown. **B)** Wild-type cells and the indicated mutants were grown in YPD media for 3 d at 30°C (YPD) and the colony grown was washed (fMAPK). The same strains were also grown in YPD supplemented with 1M KCl for 3 d at 30°C (HOG). **C)** Same cells described in panel 5A were grown in YPD (-) or YPD supplemented with 1M KCL (KCl) for 5 min at 30°C. Immunoblots were probed with p38 antibodies to detect P∼Hog1p, α-Hog1p and α-Pgk1p antibodies. P∼Hog1/Pgk1 ratio refers to relative levels of P∼Hog1p to Pgk1p of the blot shown. **D)** Same cells described in panel 5A were grown in YPD (-) or YPD supplemented with 6 µM of α-factor (α) for 5 min at 30°C. Immunoblots were probed with p44/42 antibodies to detect P∼Kss1p and P∼Fus3p, α-Kss1p, α-Fus3p and α-Pgk1p antibodies. P∼Kss1/Pgk1 ratio refers to relative levels of P∼Kss1p to Pgk1p of the blot shown. P∼Fus3/Pgk1 ratio refers to relative levels of P∼Fus3p to Pgk1p of the blot shown. **E)** Halo formation in response to α-factor of same cells described in panel 5A. Cells were spread on YPD media and α-factor was spotted at two concentrations on the surface, 2 and 6 μM, to study cell-cycle arrest. **F)** Yeast two-hybrid analysis between Cdc42p and Bem4p in wild-type cells and cells where the gene expression of *RDI1*controlled by the constitutive promoter *TEF2*.

### The polarity scaffold Bem1p interacts with the tetraspan sensor Sho1p

Bem1p is a Cdc42p-interacting protein and polarity scaffold that also regulates the mating (Leeuw et al., 1995), HOG (Tanaka et al., 2014), and fMAPK pathways (Basu et al., 2020) (**Fig. 3A**). At the polarization site and in signaling pathways, Bem1p interacts with Ste20p to promote recruitment of the protein to the plasma membrane (Winters and Pryciak, 2005). How Bem1p is itself directed to the plasma membrane in certain contexts is an open question. To begin to address this question for the fMAPK pathway, we tested for an interaction between Bem1p and the plasma membrane tetraspan protein, Sho1p. The cytosolic domain of Sho1p was expressed and purified as a His6x fusion protein from *Escherichia coli* and probed against lysates of *bem1*Δ mutant expressing pBEM1-12XMYC (provided by Daniel Lew) by a high copy (two micron) plasmid. Beads bound to the cytosolic C-terminal domain of Sho1p precipitated Bem1p-Myc from yeast extracts (**Fig. 6A**). As a control, beads bound by His6X-Sho1^CYT^ also precipitated an HA-tagged version of Pbs2p (**Fig. 6B**), which is a known Sho1p-interacting protein that functions in the HOG pathway (Maeda et al., 1995). These results identify a connection between a sensor that regulates the fMAPK pathway and a cytosolic adaptor that regulates the Cdc42p module.

**Figure 6.**
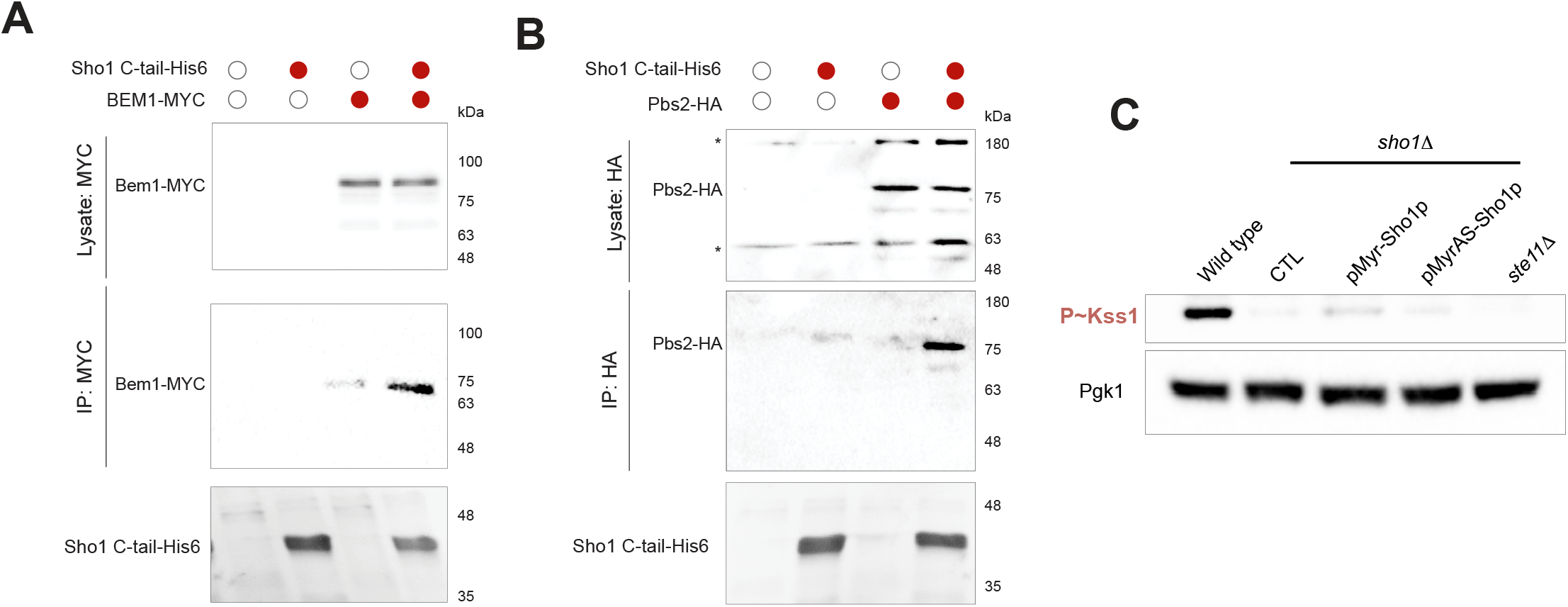
The polarity scaffold Bem1p interacts with the sensor Sho1p. **A)** The purified domain of Sho1 C-tail-His6 (SHO1^Δ*1-145*^-His6; PC3102) was probed against yeast extracts of cells expressing pRS316 or pBEM1-12XMYC (PC6512) to evaluate the interaction between the cytosolic region of Sho1p and Bem1p. **B)** The purified domain of Sho1 C-tail-His6 (SHO1^Δ*1-145*^-His6) was probed against yeast extracts of wild-type cells or containing a genomic version of *PBS2-HA* (PC1790) to evaluate the interaction between the cytosolic region of Sho1p and Bem1p. **C)** Wild-type cells, the *sho1*Δ mutant expressing a wild-type pRS316 (CTL), pMyr-Sho1p (PC4191), pMyrAS-Sho1p (PC4192), and the mutant *ste11*Δ were grown in SD-URA media for 6 h at 30°C. Immunoblots were probed with p44/42 antibodies to detect P∼Kss1p and α-Pgk1p antibodies as a loading control.

Sho1p might function solely as an adaptor to recruit proteins to the plasma membrane, or alternatively as a sensor that exists in active and inactive conformations. To test this possibility, a version of Sho1p was examined that lacks the N-terminal tetraspan domain but is anchored to the plasma membrane by an N-terminal myristoylation signal (Raitt et al., 2000). pMyr-Sho1p was unable to regulate the fMAPK pathway, based on phosphorylation Kss1p (**Fig. 6C**). pMyr-Sho1p showed the same activity as a version in which membrane anchoring was disrupted entirely (pMyrAS-Sho1). These results support the idea that Sho1p does not function only as an adaptor for the assembly of cytosolic signaling complexes but also requires domains that presumably facilitate transition of the protein to an active conformation.

## DISCUSSION

Signaling pathways regulate biological responses by protein modules that are highly conserved structurally and functionally from yeast to humans. MAPK pathways can be regulated by sensor proteins that connect to and regulate GTPase modules. By characterization of these modules, and understanding how they connect to and regulate one another, new insights into the regulation of biological processes can be appreciated. These include studies on regulated turnover, positive feedback, and the relationship between basal and activated states (English et al., 2015). Here, we report new aspects of the regulation of a Cdc42p-dependent MAPK pathway in yeast that may extend to MAPK pathway regulation in other systems.

### Interaction with Cdc42p is required for stabilization of the PAK Ste20p

A canonical aspect of MAPK pathway regulation involves the interaction with Rho GTPases, like Cdc42p, with PAK kinases (Cotteret and Chernoff, 2002). In yeast, Cdc42p interacts with the PAK Ste20p to induce the activation of MAPK pathways (Lamson et al., 2002; Leberer et al., 1997). Here we identify a new regulatory feature of PAK regulation in yeast, by turnover of the Ste20p protein in the 26S proteasome. In mammals, this regulatory feature has also been defined (Weisz Hubsman et al., 2007) and may represent an evolutionarily conserved way to modulate the activity of this type of protein kinase. In the amino-terminal domain of Ste20p, amino acid residues 333 to 370 (CRIB domain) are responsible for binding to GTP-bound Cdc42p. We found that Ste20p lacking its CRIB domain is not stable and is turned over by the 26S proteasome (**Fig. 7**). These results indicate that Cdc42p may protect Ste20p from turnover by the proteasome. This hypothesis was corroborated by examination of a version of Cdc42p that is defective for interaction with Ste20p, which also showed reduced levels of Ste20p in the cell. It is possible that Ste20p that cannot bind to Cdc42p is unstable and is rapidly turned over as part of a quality-control mechanism. In this case, Cdc42p may function to fold or otherwise stabilize Ste20p. Alternatively, Cdc42p may occlude recognition sites in the N-terminus of Ste20p by ubiquitin ligases that promote turnover of the protein. We propose a model for pathway attenuation where activated Cdc42p is degraded by proteolysis, which leads to the turnover of Ste20p, which is no longer bound to the GTPase. In this way, cells may achieve a more permanent attenuation of MAPK pathways. Further studies will be required to test this feature of Cdc42p-PAK regulation in yeast and other systems.

**Figure 7.**
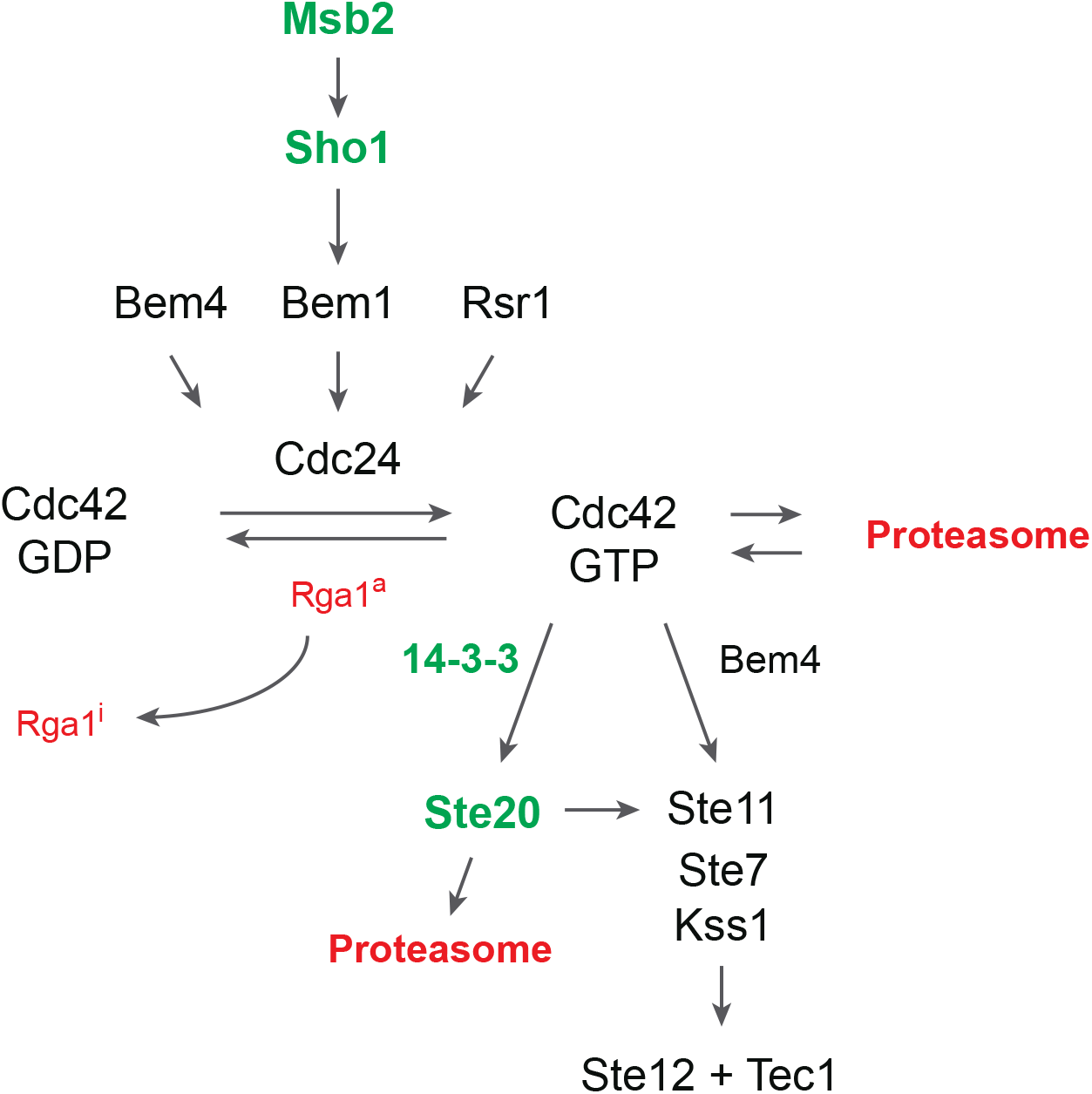
Revised model for the regulation of the fMAPK pathway. Msb2p and Sho1p sensors connect to the GTPase module through Bem1p. Negative regulatory inputs are shown in red. Ste20p is turned over in the proteosome. Rga1p inhibits basal fMAPK pathway activity (Rga1^a^, a for active) but is inhibited in some manner to prevent its inhibitory effects on an activated pathway (Rga1^i^, i for inhibited). Cdc42p induces Ste20p-dependent (green) and Ste20p-independent branches.

### A Ste20p-independent branch of the fMAPK pathway

As studies of signaling pathways become more nuanced, it is increasingly clear that pathways are not strictly linear modules but are composed of ‘branches’ that are in some cases not entirely redundant in function. One function of such branches is to allow signals to become integrated into a pathway’s activity (Yang et al., 2015). For example, RAS/PKA and TOR converge on a common set of target genes (Kunkel et al., 2019). A second example comes from the HOG pathway, which is composed of two branches (Sln1p and Ste11p) that converge on the MAPKK Pbs2p (Saito and Posas, 2012). Within the Ste11p branch, two mucins function in distinct sub-branches converging on the tetraspan protein Sho1p (Tatebayashi et al., 2007). Furthermore, Ste20p and another PAK kinase, Cla4p, constitute sub-branches of the Ste11p branch of the HOG pathway (Heinrich et al., 2007; Tatebayashi et al., 2006). While the logic of such extensive branching remains to be revealed, this knowledge helps one to understand the versatility of signaling pathways, which is probably a universal feature of signaling modules. By examining a newly characterized hyperactive version of Cdc42p, we identified a branch of the fMAPK pathway that does not require Ste20p (**Fig. 7**). This result was independently verified by examining cells lacking one of the GAPs for Cdc42p, Rga1p, which also showed Ste20p-independent activity. Like for the HOG pathway, Cdc42p might also regulate the fMAPK pathway through Cla4p. Alternatively, Cdc42p might directly activate Ste11p. In this scenario, Cdc42p would function analogously to the GTPase Ras in humans, which directly interacts with the MAPKKK, Raf. We found that the adaptor protein Bem4p, which interacts with both Cdc42p and Ste11p, was required for Ste20p-independent signaling, which might provide a connection between the two proteins. Bem4p might regulate pathway activity in cells lacking Ste20p due to its role in protecting Cdc42p; however, Cdc42p^Q61L+TD^ can bypass Bem4p in the fMAPK pathway, which identifies a potentially new function for Bem4p as a conduit between Cdc42p with Ste11p. The Ste20p-independent branch might function to increase MAPK pathway activity in certain circumstances, or sensitize the pathway to specific inputs. For example, ethanol stimulates the fMAPK pathway by a mechanism that requires Ste11p but not Ste20p (Lorenz et al., 2000).

### Cdc42p-dependent MAPK pathways function differently in basal and activating conditions

In an effort to examine the role of proteins surrounding the Cdc42p module, several common themes have emerged. First, we have confirmed that the 14-3-3 proteins regulate the fMAPK pathway, and as expected from previous findings (Roberts et al., 1997), they function in the Ste20p branch (**Fig. 7**). These proteins, at least when overexpressed, promote signaling from the basal to activated state of the pathway. Interestingly, these proteins do not hyperactivate all three Cdc42p-dependent MAPK pathways, but rather (modestly) dampen basal HOG and mating activity. This suggests that there is a pathway-specific association between these proteins and other fMAPK pathway-specific regulators, such as Bem4p. Recently Bmh1p and Bmh2p have been identified as key regulators of meiotic commitment in yeast (Gavade et al., 2022). These proteins also impact retrograde mitochondria-to-nucleus signaling (Liu et al., 2003), and other pathways such as the Target of Rapamycin or TOR pathway (Bertram et al., 1998). Given that filamentous growth occurs differently in haploids and diploids (Roberts and Fink, 1994), and is impacted by the retrograde (Jin et al., 2008) and TOR pathways (Cutler et al., 2001), the 14-3-3 proteins might impact filamentous growth and signaling at multiple levels.

We also found that Rga1p, the main GAP for Cdc42p in the fMAPK pathway, exerts its effects on Cdc42p mainly in the basal state (**Fig. 7**). How is Rga1p restricted from attenuating active MAPK pathways? Rga1p may get evicted from complexes containing active Cdc42p. It may be excluded by PAK binding to Cdc42p or modified by a post-translational modification to inactive its GAP activity. Rga1p’s localization is critical for its function (Miller et al., 2017), and it may be that its localization is altered in response to MAPK pathway activation. Rga1p is important for attenuating basal signaling, while proteasomal Cdc42p degradation may regulate activated signaling. Perhaps another GAP is responsible for attenuating activated signaling. This idea is not supported by phenotypic analysis, because other GAPs do not show phenotypes in filamentous growth. This idea is also not supported by genetic screens for negative regulators of the fMAPK pathway, which have typically identified a single GAP, Rga1p.

Our results also identify distinctions between the ways that signaling pathways are regulated in basal and activated states. We have previously shown that bud-site-selection proteins regulate the fMAPK pathway, and their roles are more critical during basal signaling (Basu et al., 2016). Rga1p may be more effective at dampening the fMAPK through a mechanism involving bud-site-selection proteins. Indeed, Rga1p is critical for preventing budding at previous bud sites (Meitinger et al., 2014; Meitinger et al., 2013; Tong et al., 2007).

We also show that there are higher levels of phosphorylated Kss1p than phosphorylated Fus3p under basal conditions. This might be because Fus3p is conformationally constrained (locked) such that it is only activated in the presence of pheromone (Good et al., 2009). These results imply that cells may be more potentiated for filamentous growth than for mating, even under nutrient-replete conditions. In line with this idea, cells under nutrient-replete conditions exhibit some fMAPK-dependent phenotypes, such as cell adhesion (Pitoniak et al., 2015).

### Exploring GEF regulation during filamentous growth

An important open question surrounding the fMAPK pathway is to understand how the sensors connect to and regulate the GTPase module. Central to this question is the GEF Cdc24p, which regulates Cdc42p activation during filamentous growth. Our results indicate that the phosphorylation of Cdc24p, or the levels of the protein are not be critical determinants for the regulation of the fMAPK pathway. Here, we show that Bem1p, a Cdc24p-interacting protein and regulator of the fMAPK pathway, associates with the tetraspan protein Sho1p (**Fig. 7**). Interestingly, Bem1p also associates with the signaling mucin Msb2p in the HOG pathway (Tanaka et al., 2014). In addition to Bem1p, Sho1p associates with Ste20p and Ste11p (Tatebayashi et al., 2006), and is considered to function as an adaptor protein, bringing multiple signaling proteins together. We cannot exclude the possibility that Sho1p associates with Bem1p indirectly through its interaction with Ste20p, given that Ste20p and Bem1p directly associate (Winters and Pryciak, 2005). In any event, our results connect a sensor for the fMAPK that functions at the plasma membrane to a regulator of the Cdc42p module. Perhaps this connection will begin to provide clues for how GEF activation occurs in a mucin-dependent signaling pathway.

## MATERIALS AND METHODS

### Yeast strains, reagents and media

Strains are listed in Table 1, and plasmids are listed in Table 2. Yeast were grown in synthetic media (SD; 0.67% yeast nitrogen base without amino acids, 2% dextrose), supplemented with amino acids as required, <u>Y</u>east <u>E</u>xtract <u>P</u>eptone <u>D</u>extrose media [YEPD (1% bacto-yeast extract, 2% bacto-peptone, 2% dextrose)] and YEPGAL (1% bacto-yeast extract, 2% bacto-peptone, 2% galactose) at 30°C. Gene disruptions were performed by antibiotic resistance markers *NAT*, *HYG* and *KanMX6* using PCR-based approaches using published templates (Goldstein and McCusker, 1999; Longtine et al., 1998). Yeast strains and plasmids have been previously described. Some gene deletions were constructed using cassettes that contained antibiotic resistance markers. Plasmid pRL116 CEN/URA4 pGFP-Ste20p and pGFP-Ste20p-CRIBΔ were provided by P. Pryciak (Leberer et al., 1997). Plasmids pRS315 and pRS316 have been described (Sikorski and Hieter, 1989). The Cdc24p phosphorylation sites wild type (PC3622) and mutants including Cdc24^PH-A^ (PC3624), Cdc24^Linker-A^ (PC3625), Cdc24^PH-A, Linker-A^ (PC3623), and Cdc24^T35A^ (PC3626) were generously shared by Rong Li (Wai et al., 2007). Plasmids carrying Bem1p-12XMYC (pDLB2374) were provided by Daniel Lew (Irazoqui et al., 2003). Two-hybrid analysis was performed as described in (Basu et al., 2020). Tests for viability and function of *CDC24* alleles were performed by plasmid loss experiments of the pCDC24 based URA3 plasmid on 5-FOA (Boeke et al., 1984). pCDC24-lacZ (PC2107) was constructed by *in vivo* recombination of the PCR-amplified *CDC24* gene into vector V84 provided by C. Boone.

**Table 1.**
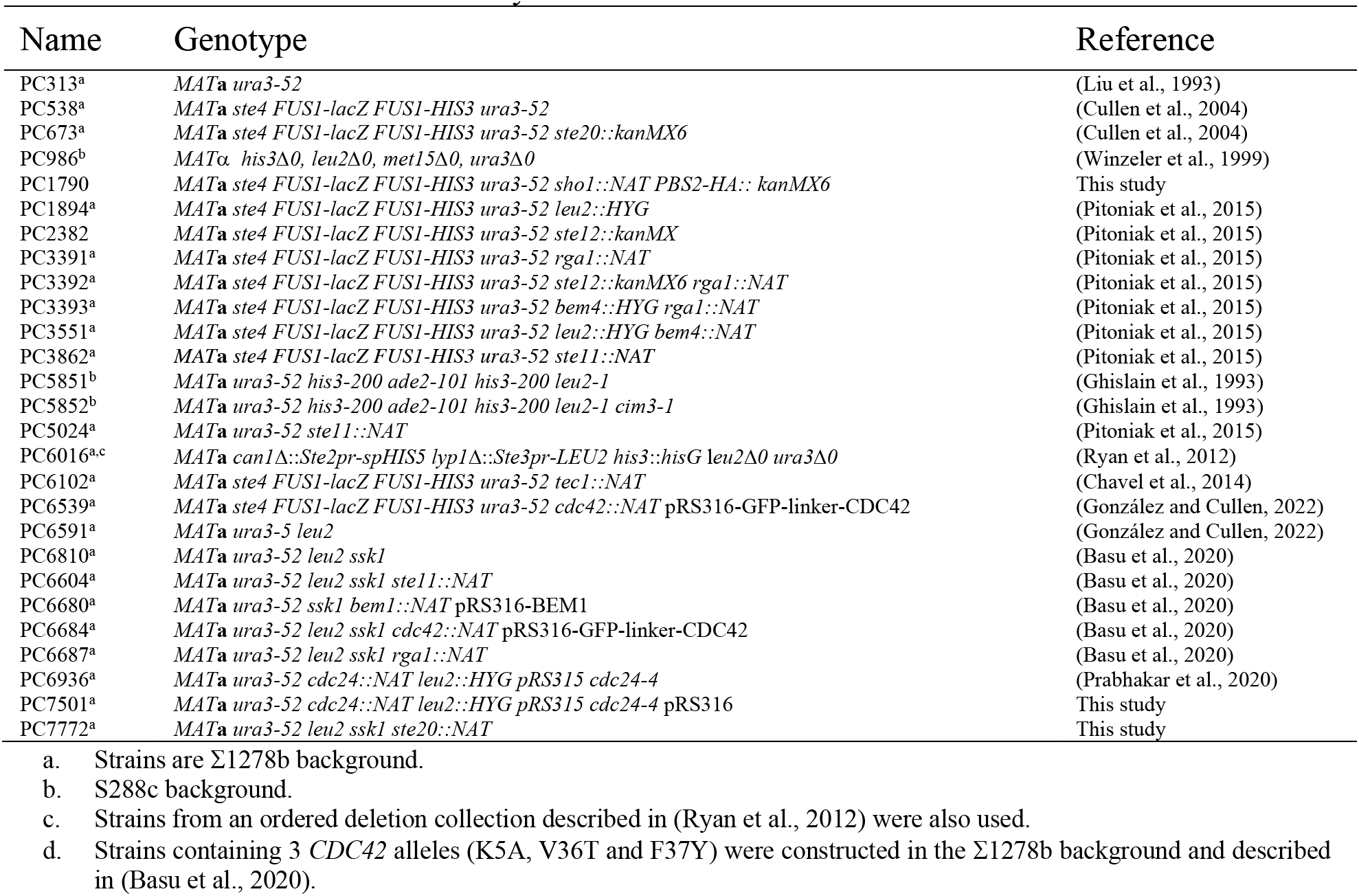
Yeast strains used in the study.

**Table 2.**
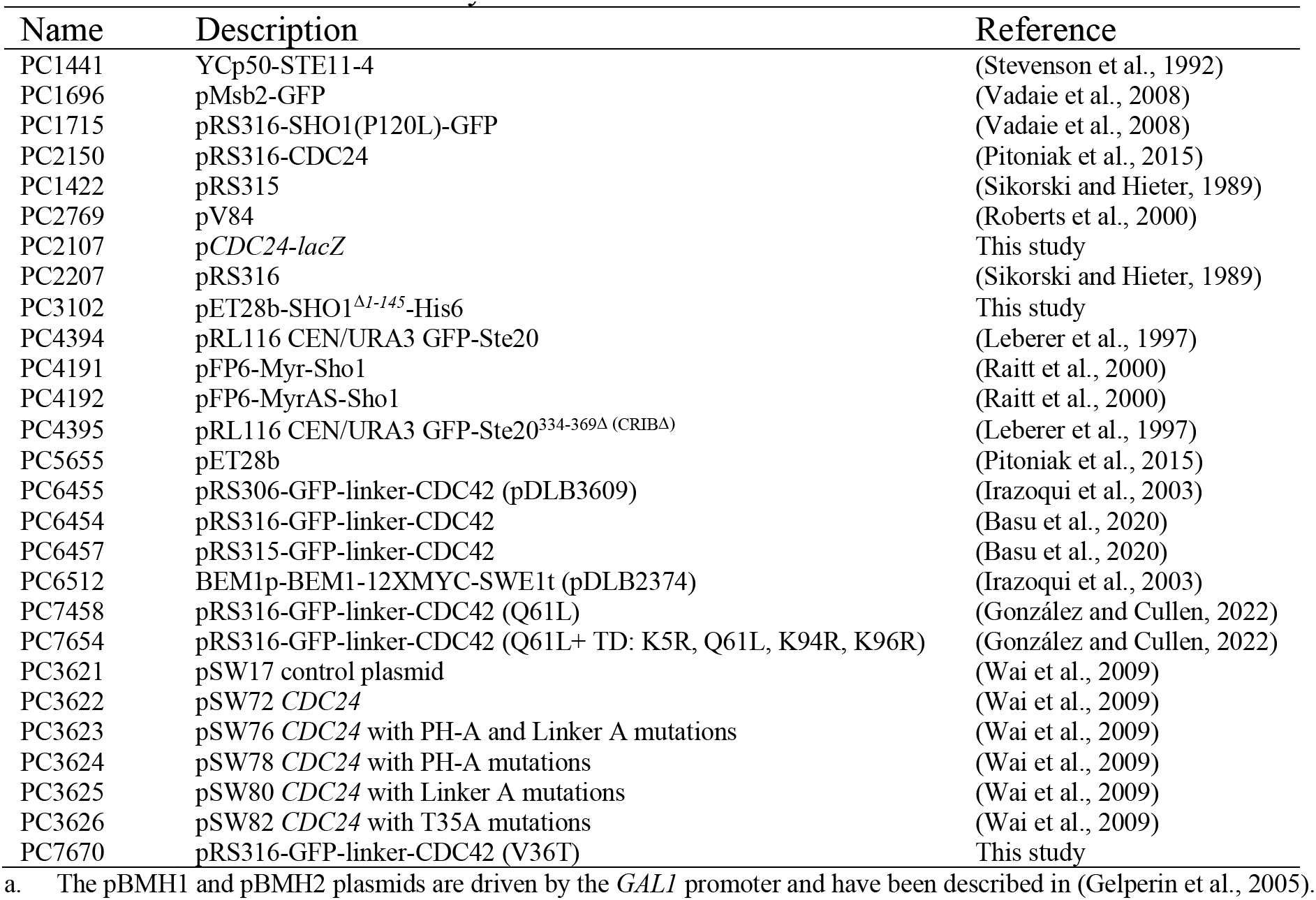
Plasmids used in the study.

### In vitro pulldowns

The interaction between Sho1p^D1-145^-His6 (pSho1-C-tail-His6; PC3102) and Bem1p was tested by the following method. 250 ml of *E. coli* containing pET28b-Sho1-C-tail-His6 or pET28b were grown to exponential phase in 2XYT broth containing kanamycin and induced with 10 µM of IPTG (isopropyl-b-d-1-thiogalactopyranoside) for 5 h at 30°C. Cells were centrifuged at 4ªC at 12,000 rpm and frozen at -80°C. For protein Sho1p^D*1-145*^-His6 purification, 10 µL/ml of EDTA-Free Protease Inhibitor Cocktail (Cat# 4693124001, Roche), and 20 ml of Tractor Buffer (Cat# 63567, TAKARA), followed by 2.5U/ml of DNase I and 50 mg/ml of lysozyme were added and incubated for 10 min at 25°C. Cell debris were removed by centrifugation at 20,000 rpm for 20 min at 4°C. The supernatant was applied to 250 µL Talon metal affinity resin (Cat# 635652, TAKARA) previously washed twice with Equilibration buffer, and incubated with rotation for 20 min at 4°C. The resin was then washed three times with Equilibration buffer and applied to a 10 mL of gravity-flow column and washed twice with 5 bed volumes of Washing Buffer. 50 mL of *S. cerevisiae* lacking Bem1p and expressing the pBEM1-12XMYC plasmid were grown to mid-log phase in SD-LEU media. Yeast extracts were obtained by centrifugation, and washed twice with 1% of PBS (phosphate-buffered saline; 137 mM NaCl, 2.7 mM KCl, 10 mM Na_2_HPO_4_, 1.8 mM KH_2_PO_4_) and resuspended in 500 µl of Immuno precipitation buffer (50 mM Tris-Cl, pH 8, 1 mM EDTA, 100 mM NaCl, 1.5% NP-40, 1 mM phenylmethanesulfonyl fluoride (PMSF), 1x protease inhibitor cocktail). Then, the lysed cells were centrifuged at 4°C for 10 min at 14,000 rpm and the supernatant was applied to the gravity-flow column containing the purified Sho1p^D*1-145*^-His6. The column was washed once with washing buffer, and eluted with 1 ml of Elution Buffer (150 mM imidazole, 50 mM sodium phosphate [pH 7.0], 100 mM NaCl). Proteins were evaluated by SDS-PAGE analysis.

### Immunoblot and Phosphoblot Analysis

Cells were grown to saturation in SD or YEPD media for 16 h and transferred fresh media and grown for 4-6 h to mid log phase. Cells were harvested by centrifugation. Proteins extracts were prepared by mechanical disruption with beads followed by a trichloroacetic acid (TCA) precipitation method (Basu et al., 2020). Protein precipitates were analyzed by SDS-PAGE and transferred to a nitrocellulose membrane (Cat#10600003, Amersham^TM^ Protran^TM^ Premium 0.45 μm NC, GE Healthcare Life sciences). Monoclonal mouse anti-GFP antibodies were used (Cat#11814460001, clones 7.1 and 13.1, Roche) at 1:1,000 dilution. Polyclonal rabbit phospho-p44/42 MAPK (Erk1/2, Cat#3102, Cell Signaling Technology), and p38 (Cat#9211, Cell Signaling Technology) were used at 1:10,000 dilution. Mouse anti-Kss1p antibodies (yC-19, Santa Cruz Biotechnology) and anti-Hog1p antibodies (Cat# yC-20, Santa Cruz Biotechnology) were used at 1:10,000 dilution. Monoclonal mouse anti-Pgk1p antibodies (22C5D8, Cat#459250, Invitrogen) were 1:1,000 dilution. Secondary anti-mouse IgG-HRP (Cat# 1706516, Bio-Rad Laboratories) and goat anti-rabbit IgG-HRP (Cat#115-035-003, Jackson ImmnunoResearch Laboratories) were used. The nitrocellulose membrane was blocked with 5% non-fat dried milk or 5% bovine serum albumin (BSA) (BSA; for p44/42 antibody) for 1 h prior antibody detection. Primary incubations were performed at 4°C for 16 h and secondary at 20°C for 1 h. IBs were visualized by Gel Doc XR Imaging System (Bio-Rad, Inc.), after addition of Chemiluminescent HRP substrate for chemiluminescent Westerns (Radiance^TM^ Plus Substrate, Azure Biosystems).

Band intensities quantitation of P∼Fus3p, P∼Kss1p, Cdc42p, GFP-Cdc42p and ubiquitin were detected under non-saturated conditions and normalized to the housekeeping protein Pgk1p using the Image Lab Software (Bio-Rad, Inc.). Wild-type cells and control conditions were set to a value of one and adjusted for other samples accordingly.

### MAPK pathway reporters and functional tests

The fMAPK pathway activity was evaluated by assessing the *FUS1-HIS3* growth reporter in cells lacking *STE4* (Adhikari et al., 2015; McCaffrey et al., 1987). Cells lacking an intact mating pathway (*ste4*Δ) mating, show basal activity of the fMAPK pathway by this reporter (Cullen et al., 2004). Wild-type cells (PC538) and a control strain (*ste11*Δ, PC3862), were growth in SD-URA media to maintain plasmid selection (SD-URA-HIS, control), and media lacking Histidine (SD-URA-HIS) or supplemented with ATA (3-amino-1,2,4-triazole) to evaluate fMAPK activity.

The *FRE-lacZ* (Madhani and Fink, 1997) and *FUS1-LacZ* (*ste4* background) (Chavel et al., 2014) reporters were also used to study the fMAPK pathway activity. The plate washing was performed as described in (Roberts and Fink, 1994) to evaluate the fMAPK pathway phenotype. To evaluate the HOG pathway, cells were grown in media supplemented with 0.5 M KCl or 1 M KCl and analyzed by phophoblot analysis or by growth on semi-solid agar media. The mating pathway activity was also evaluated by phosphoblot analysis, halo assays, and shmoo formation by microscopy. For halo assays, cells were spread on SD-URA or YPD semi-solid agar media. After the liquid had evaporated, alpha-factor was applied to the surface of the plate, which were incubated for 2 d at 30°C.

### Fluorescence microscopy

Differential interference contrast (DIC), fluorescence microscopy using fluorescein isothiocyanate (FITC) and Rhodamine filter sets were used in an Axioplan 2 fluorescence microscope (Zeiss) with a Paln-Apochromat 100x/1.4 (oil) objective with the Axiocam MRm camera (Zeiss). Images were analyzed using Axiovision 4.4 software (Zeiss). Actin stanning was performed as described (Amberg et al., 2006) using Phalloidin-Atto 532 (Millpore Sigma, MA, 49429). Images were analyzed in Adobe Photoshop and ImageJ. Fluorescence images were converted to Grayscale and inverted using ImageJ.

### Statistical analysis

All statistical tests were performed with Prism 7 (GraphPad; https://www.graphpad.com/scientific-software/prism). Data were analyzed by one-way ANOVA test followed by a Tukey’s multiple comparison test to generate p-values.

## Data availability statement

All the data are included in the manuscript and supplemental files and are available upon request at.

## ABBREVIATIONS

ATA: 3-amino-1,2,4 triazole
5-FOA: 5-floroorotic acid
CRIB: Rac-interactive binding motif
GAP: GTPase activating protein
GEF: guanine nucleotide exchange factor
GDI: guanine nucleotide dissociation inhibitor
HIS: histidine
MAPK: mitogen-activated protein kinase
PM: plasma membrane
PAK: p21-activated kinase
SD: standard deviation
URA: uracil

## ACKNOWLEDGEMENTS

Thanks to Charlie Boone (University of Toronto), Daniel Lew (Duke University), Rong Li (Johns Hopkins University), Haruo Saito (University of Tokyo), and Peter Pryciak (UMASS) for providing plasmids. Thanks to Nadia Vadaie, Colin Chavel, and Heather Dionne for assistance with experiments. Thanks to Matt Vandermeulen and other members of the lab for suggestions. The work was supported by a grant from the NIH (GM098629).

## SUPPORTING INFORMATION

**Figure S1. Evaluating signaling by an activated version of Cdc42p in the fMAPK, HOG and mating pathways and the indicated mutants. A)** Wild-type cells (WT, PC538) and the *ste11*Δ (PC3862), *ste20*Δ (PC673) and *ste12*Δ (PC2382) mutants expressing pGFP-Cdc42p (Cdc42; PC6454), pGFP-Cdc42p^Q61L^ (Q61L; PC7458) or pGFP-Cdc42p^Q61L+TD^ (Q61L+TD: PC7654) were grown on SD-URA, SD-URA-HIS or SD-URA-HIS supplemented with 3-amino-1,2,4,-triazole (ATA) media to evaluate the activity of the *FUS1-HIS3* reporter. **B)** Cells lacking Ste20p and expressing pGFP-Cdc42p, pGFP-Cdc42p^Q61L^ or pGFP-Cdc42p^Q61L+TD^ were grown in YPD for 10 days. Colonies were washed in a stream of water and the plate was photographed. **C)** Wild-type cells (WT, PC6810), and the *ste20*Δ (PC7772), the *ste11*Δ (PC6604) mutants in the *ssk1*Δ background expressing plasmids described in panel S1A were grown on YPGAL (Gal) or YGAL supplemented with 0.5 M KCl (GAL + 0.5 M KCl). **D)** Same cells described in panel S1B were spread on SD-URA media and α-factor was top added at two concentrations, 6 and 2 µM, to study cell cycle arrest. **E)** Wild-type cells, and the s*te20*Δ and *ste11*Δ mutans expressing pRS316 (PC2207), Msb2-GFP (PC1696), Sho1pP120L (PC1715), Cdc42p^Q61L^, Cdc42p^Q61L+TD^, or Ste11-4 (PC1441) were grown in SGAL-URA media to evaluate the activity of the *FUS1-HIS3* reporter.

**Fig. S2. The 14-3-3 proteins Bmh1p and Bmh2p regulate the fMAPK pathway. A)** Wild-type cells (PC538, *ste4*) and several independent isolates lacking Bmh1p or Bmh2p as indicated were spotted on SD+AA and SD-URA media to evaluate the activity of the *FUS1-HIS3* reporter. **B)** Cells described in panel 3D (GAL) and 3F (GAL+ α-factor) were observed by microscopy at the 3 h. Scale bar, 10 µm.

**Figure S3. Role of phosphorylation of Cdc24p in regulating the fMAPK pathway. A)** Cells lacking Cdc24p and expressing p*CDC24* and p*CDC24-4* were grown on SD and SD-HIS media to study the fMAPK pathway activity. **B)** Serial dilutions of cells lacking Cdc24p and expressing Cdc24^PH-A^, Cdc24^Linker-A^, Cdc24^PH-A, Linker-A^, or Cdc24^T35A^ [provided by Rong Li (Wai et al., 2009) were spotted on SD-URA-LEU, SD-LEU +FOA. Cells grown on the FOA plate were grown on SD-LEU, and SD-LEU-HIS+10ul ATA media (2.5M). Cells were grown for 2 days at 30°C. **C)** Same cells described in panel 6B and the *ste11*Δ mutant were grown in YEPD to mid-log phase + 6 hours in YEP-GAL to induce filamentous growth. Cell extracts were probed for phosphorylated Kss1p (P∼Kss1p), Kss1p, and Pgk1p by immunoblotting using anti-phospho p44/42, Kss1p, and Pgk1p antibodies. **D)** Cell extracts derived from cells expressing p*CDC24-lacZ* were examined for ß-galactosidase activity, expressed as Miller Units in independent biological replicates. The difference between wild-type cells and the *ste12* mutant was not statistically significant.

## REFERENCES

Adhikari, H., and P.J. Cullen. 2014. Metabolic Respiration Induces AMPK- and Ire1p-Dependent Activation of the p38-Type HOG MAPK Pathway. PLOS Genetics. 10:e1004734.

Adhikari, H., N. Vadaie, J. Chow, L.M. Caccamise, C.A. Chavel, B. Li, A. Bowitch, C.J. Stefan, and P.J. Cullen. 2015. Role of the unfolded protein response in regulating the mucin-dependent filamentous-growth mitogen-activated protein kinase pathway. Mol Cell Biol. 35:1414–1432.

Amberg, D.C., D.J. Burke, and J.N. Strathern. 2006. Actin staining in fixed yeast cells. CSH Protoc. 2006.

Ash, J., C. Wu, R., Larocque, M., Jamal, W., Stevens, M. Osborne, D.Y. Thomas, and M. Whiteway. 2003. Genetic analysis of the interface between Cdc42p and the CRIB domain of Ste20p in Saccharomyces cerevisiae. Genetics. 163:9–20.

Bao, M.Z., M.A. Schwartz, G.T. Cantin, J.R. Yates, 3rd, and H.D. Madhani. 2004. Pheromone-dependent destruction of the Tec1 transcription factor is required for MAP kinase signaling specificity in yeast. Cell. 119:991–1000.

Bardwell, L. 2005. A walk-through of the yeast mating pheromone response pathway. Peptides. 26:339–350.

Bardwell, L., J.G. Cook, J.X. Zhu-Shimoni, D. Voora, and J. Thorner. 1998. Differential regulation of transcription: Repression by unactivated mitogen-activated protein kinase Kss1 requires the Dig1 and Dig2 proteins. Proceedings of the National Academy of Sciences. 95:15400–15405.

Bassilana, M., J. Blyth, and R.A. Arkowitz. 2003. Cdc24, the GDP-GTP exchange factor for Cdc42, is required for invasive hyphal growth of Candida albicans. Eukaryot Cell. 2:9–18.

Basu, S., B. González, B. Li, G. Kimble, K.G. Kozminski, and P.J. Cullen. 2020. Functions for Cdc42p BEM adaptors in regulating a differentiation-type MAP kinase pathway. Mol Biol Cell. 31:491–510.

Basu, S., N. Vadaie, A. Prabhakar, B. Li, H. Adhikari, A. Pitoniak, J. Chow, C.A. Chavel, and P.J. Cullen. 2016. Spatial landmarks regulate a Cdc42-dependent MAPK pathway to control differentiation and the response to positional compromise. Proceedings of the National Academy of Sciences. 113:E2019.

Bertram, P.G., C. Zeng, J. Thorson, A.S. Shaw, and X.F. Zheng. 1998. The 14-3-3 proteins positively regulate rapamycin-sensitive signaling. Current biology : CB. 8:1259–1267.

Boeke, J.D., F. LaCroute, and G.R. Fink. 1984. A positive selection for mutants lacking orotidine-5’-phosphate decarboxylase activity in yeast: 5-fluoro-orotic acid resistance. Mol Gen Genet. 197:345–346.

Bose, I., J.E. Irazoqui, J.J. Moskow, E.S. Bardes, T.R. Zyla, and D.J. Lew. 2001. Assembly of scaffold-mediated complexes containing Cdc42p, the exchange factor Cdc24p, and the effector Cla4p required for cell cycle-regulated phosphorylation of Cdc24p. Journal of Biological Chemistry. 276:7176–7186.

Boulter, E., R. Garcia-Mata, C. Guilluy, A. Dubash, G. Rossi, P.J. Brennwald, and K. Burridge. 2010. Regulation of Rho GTPase crosstalk, degradation and activity by RhoGDI1. Nat Cell Biol. 12:477–483.

Braicu, C., M. Buse, C. Busuioc, R. Drula, D. Gulei, L. Raduly, A. Rusu, A. Irimie, A.G. Atanasov, O. Slaby, C. Ionescu, and I. Berindan-Neagoe. 2019. A Comprehensive Review on MAPK: A Promising Therapeutic Target in Cancer. Cancers (Basel). 11.

Chavel, C.A., L.M. Caccamise, B. Li, and P.J. Cullen. 2014. Global regulation of a differentiation MAPK pathway in yeast. Genetics. 198:1309–1328.

Chou, S., L. Huang, and H. Liu. 2004. Fus3-regulated Tec1 degradation through SCFCdc4 determines MAPK signaling specificity during mating in yeast. Cell. 119:981–990.

Chow, J., I. Starr, S. Jamalzadeh, O. Muniz, A. Kumar, O. Gokcumen, D.M. Ferkey, and P.J. Cullen. 2019. Filamentation Regulatory Pathways Control Adhesion-Dependent Surface Responses in Yeast. Genetics. 212:667–690.

Cook, J.G., L. Bardwell, S.J. Kron, and J. Thorner. 1996. Two novel targets of the MAP kinase Kss1 are negative regulators of invasive growth in the yeast Saccharomyces cerevisiae. Genes Dev. 10:2831–2848.

Cook, J.G., L. Bardwell, and J. Thorner. 1997. Inhibitory and activating functions for MAPK Kss1 in the S. cerevisiae filamentous-growth signalling pathway. Nature. 390:85–ΔΔΔΔΔ.

Cotteret, S., and J. Chernoff. 2002. The evolutionary history of effectors downstream of Cdc42 and Rac. Genome Biology. 3:reviews0002.0001.

Cullen, P.J., W. Sabbagh, Jr., E. Graham, M.M. Irick, E.K. van Olden, C. Neal, J. Delrow, L. Bardwell, and G.F. Sprague, Jr. 2004. A signaling mucin at the head of the Cdc42- and MAPK-dependent filamentous growth pathway in yeast. Genes Dev. 18:1695–1708.

Cutler, N.S., X. Pan, J. Heitman, and M.E. Cardenas. 2001. The TOR signal transduction cascade controls cellular differentiation in response to nutrients. Mol Biol Cell. 12:4103–4113.

English, J.G., J.P. Shellhammer, M. Malahe, P.C. McCarter, T.C. Elston, and H.G. Dohlman. 2015. MAPK feedback encodes a switch and timer for tunable stress adaptation in yeast. Sci Signal. 8:ra5.

Fey, D., D. Croucher, W. Kolch, and B. Kholodenko. 2012. Crosstalk and Signaling Switches in Mitogen-Activated Protein Kinase Cascades. Front Physiol. 3.

Gavade, J.N., C.M. Puccia, S.G. Herod, J.C. Trinidad, L.E. Berchowitz, and S. Lacefield. 2022. Identification of 14-3-3 proteins, Polo kinase, and RNA-binding protein Pes4 as key regulators of meiotic commitment in budding yeast. Current Biology. 32:1534–1547.e1539.

Gelperin, D.M., M.A. White, M.L. Wilkinson, Y. Kon, L.A. Kung, K.J. Wise, N. Lopez-Hoyo, L. Jiang, S. Piccirillo, H. Yu, M. Gerstein, M.E. Dumont, E.M. Phizicky, M. Snyder, and E.J. Grayhack. 2005. Biochemical and genetic analysis of the yeast proteome with a movable ORF collection. Genes Dev. 19:2816–2826.

Ghislain, M., A. Udvardy, and C. Mann. 1993. S. cerevisiae 26S protease mutants arrest cell division in G2/metaphase. Nature. 366:358–362.

Gimeno, C.J., P.O. Ljungdahl, C.A. Styles, and G.R. Fink. 1992. Unipolar cell divisions in the yeast S. cerevisiae lead to filamentous growth: regulation by starvation and RAS. Cell. 68:1077–1090.

Gladfelter, A.S., J.J. Moskow, T.R. Zyla, and D.J. Lew. 2001. Isolation and characterization of effector-loop mutants of CDC42 in yeast. Mol Biol Cell. 12:1239–1255.

Golding, A.E., I. Visco, P. Bieling, and W.M. Bement. 2019. Extraction of active RhoGTPases by RhoGDI regulates spatiotemporal patterning of RhoGTPases. eLife. 8:e50471.

Goldstein, A.L., and J.H. McCusker. 1999. Three new dominant drug resistance cassettes for gene disruption in Saccharomyces cerevisiae. Yeast. 15:1541–1553.

González, B., and P.J. Cullen. 2022. Regulation of Cdc42 protein turnover modulates the filamentous growth MAPK pathway. Journal of Cell Biology. 221.

Good, M., G. Tang, J. Singleton, A. Reményi, and W.A. Lim. 2009. The Ste5 scaffold directs mating signaling by catalytically unlocking the Fus3 MAP kinase for activation. Cell. 136:1085–1097.

Grice, G.L., and J.A. Nathan. 2016. The recognition of ubiquitinated proteins by the proteasome. Cell Mol Life Sci. 73:3497–3506.

Gulli, M.-P., M. Jaquenoud, Y. Shimada, G. Niederhäuser, P. Wiget, and M. Peter. 2000. Phosphorylation of the Cdc42 exchange factor Cdc24 by the PAK-like kinase Cla4 may regulate polarized growth in yeast. Molecular cell. 6:1155–1167.

Hao, N., N. Yildirim, M.J. Nagiec, S.C. Parnell, B. Errede, H.G. Dohlman, and T.C. Elston. 2012. Combined computational and experimental analysis reveals mitogen-activated protein kinase-mediated feedback phosphorylation as a mechanism for signaling specificity. Mol Biol Cell. 23:3899–3910.

Heinrich, M., T. Köhler, and H.U. Mösch. 2007. Role of Cdc42-Cla4 interaction in the pheromone response of Saccharomyces cerevisiae. Eukaryot Cell. 6:317–327.

Irazoqui, J.E., A.S. Gladfelter, and D.J. Lew. 2003. Scaffold-mediated symmetry breaking by Cdc42p. Nat Cell Biol. 5:1062–1070.

Jamalzadeh, S., A.N. Pujari, and P.J. Cullen. 2020. A Rab escort protein regulates the MAPK pathway that controls filamentous growth in yeast. Sci Rep. 10:22184.

Jin, R., C.J. Dobry, P.J. McCown, and A. Kumar. 2008. Large-scale analysis of yeast filamentous growth by systematic gene disruption and overexpression. Mol Biol Cell. 19:284–296.

Johnson, D.I. 1999. Cdc42: An essential Rho-type GTPase controlling eukaryotic cell polarity. Microbiology and molecular biology reviews : MMBR. 63:54–105.

Karunanithi, S., and P.J. Cullen. 2012. The filamentous growth MAPK Pathway Responds to Glucose Starvation Through the Mig1/2 transcriptional repressors in Saccharomyces cerevisiae. Genetics. 192:869–887.

Kim, J., and M.D. Rose. 2022. Cla4p Kinase Activity Is Down-Regulated by Fus3p during Yeast Mating. Biomolecules. 12.

Kolch, W. 2005. Coordinating ERK/MAPK signalling through scaffolds and inhibitors. Nat Rev Mol Cell Biol. 6:827–837.

Kozminski, K.G., A.J. Chen, A.A. Rodal, and D.G. Drubin. 2000. Functions and functional domains of the GTPase Cdc42p. Molecular biology of the cell. 11:339–354.

Kunkel, J., X. Luo, and A.P. Capaldi. 2019. Integrated TORC1 and PKA signaling control the temporal activation of glucose-induced gene expression in yeast. Nature Communications. 10:3558.

Lamson, R.E., M.J. Winters, and P.M. Pryciak. 2002. Cdc42 regulation of kinase activity and signaling by the yeast p21-activated kinase Ste20. Molecular and cellular biology. 22:2939–2951.

Leberer, E., C. Wu, T. Leeuw, A. Fourest-Lieuvin, J.E. Segall, and D.Y. Thomas. 1997. Functional characterization of the Cdc42p binding domain of yeast Ste20p protein kinase. EMBO J. 16:83–97.

Leeuw, T., A. Fourest-Lieuvin, C. Wu, J. Chenevert, K. Clark, M. Whiteway, D.Y. Thomas, and E. Leberer. 1995. Pheromone response in yeast: association of Bem1p with proteins of the MAP kinase cascade and actin. Science. 270:1210–1213.

Leeuw, T., C. Wu, J.D. Schrag, M. Whiteway, D.Y. Thomas, and E. Leberer. 1998. Interaction of a G-protein beta-subunit with a conserved sequence in Ste20/PAK family protein kinases. Nature. 391:191–195.

Liu, H., C.A. Styles, and G.R. Fink. 1993. Elements of the yeast pheromone response pathway required for filamentous growth of diploids. Science. 262:1741–1744.

Liu, Z., T. Sekito, M. Spírek, J. Thornton, and R.A. Butow. 2003. Retrograde signaling is regulated by the dynamic interaction between Rtg2p and Mks1p. Mol Cell. 12:401–411.

Longtine, M.S., A. McKenzie, 3rd, D.J. Demarini, N.G. Shah, A. Wach, A. Brachat, P. Philippsen, and J.R. Pringle. 1998. Additional modules for versatile and economical PCR-based gene deletion and modification in Saccharomyces cerevisiae. Yeast. 14:953–961.

Lorenz, M.C., N.S. Cutler, and J. Heitman. 2000. Characterization of alcohol-induced filamentous growth in Saccharomyces cerevisiae. Mol Biol Cell. 11:183–199.

Lyons, D.M., S.K. Mahanty, K.Y. Choi, M. Manandhar, and E.A. Elion. 1996. The SH3-domain protein Bem1 coordinates mitogen-activated protein kinase cascade activation with cell cycle control in Saccharomyces cerevisiae. Mol Cell Biol. 16:4095–4106.

Madhani, H.D., and G.R. Fink. 1997. Combinatorial control required for the specificity of yeast MAPK signaling. Science. 275:1314–1317.

Madhani, H.D., T. Galitski, E.S. Lander, and G.R. Fink. 1999. Effectors of a developmental mitogen-activated protein kinase cascade revealed by expression signatures of signaling mutants. Proceedings of the National Academy of Sciences. 96:12530.

Maeda, T., M. Takekawa, and H. Saito. 1995. Activation of yeast PBS2 MAPKK by MAPKKKs or by binding of an SH3-containing osmosensor. Science. 269:554–558.

Malleshaiah, M.K., V. Shahrezaei, P.S. Swain, and S.W. Michnick. 2010. The scaffold protein Ste5 directly controls a switch-like mating decision in yeast. Nature. 465:101–105.

McCaffrey, G., F.J. Clay, K. Kelsay, and G.F. Sprague, Jr. 1987. Identification and regulation of a gene required for cell fusion during mating of the yeast Saccharomyces cerevisiae. Mol Cell Biol. 7:2680–2690.

McCullagh, E., A. Seshan, H. El-Samad, and H.D. Madhani. 2010. Coordinate control of gene expression noise and interchromosomal interactions in a MAP kinase pathway. Nat Cell Biol. 12:954–962.

Meitinger, F., A. Khmelinskii, S. Morlot, B. Kurtulmus, S. Palani, A. Andres-Pons, B. Hub, M. Knop, G. Charvin, and G. Pereira. 2014. A memory system of negative polarity cues prevents replicative aging. Cell. 159:1056–1069.

Meitinger, F., H. Richter, S. Heisel, B. Hub, W. Seufert, and G. Pereira. 2013. A safeguard mechanism regulates Rho GTPases to coordinate cytokinesis with the establishment of cell polarity. PLoS Biol. 11:e1001495.

Miller, K.E., W.C. Lo, M.E. Lee, P.J. Kang, and H.O. Park. 2017. Fine-tuning the orientation of the polarity axis by Rga1, a Cdc42 GTPase-activating protein. Mol Biol Cell. 28:3773–3788.

Min, K., A.M. Neiman, and J.B. Konopka. 2020. Fungal Pathogens: Shape-Shifting Invaders. Trends Microbiol. 28:922–933.

Mitchell, A.P. 1998. Dimorphism and virulence in Candida albicans. Curr Opin Microbiol. 1:687–692.

Moran, K.D., H. Kang, A.V. Araujo, T.R. Zyla, K. Saito, D. Tsygankov, and D.J. Lew. 2019. Cell-cycle control of cell polarity in yeast. J Cell Biol. 218:171–189.

Morrison, D.K. 2012. MAP kinase pathways. Cold Spring Harbor perspectives in biology. 4:a011254.

Moskow, J.J., A.S. Gladfelter, R.E. Lamson, P.M. Pryciak, and D.J. Lew. 2000. Role of Cdc42p in pheromone-stimulated signal transduction in Saccharomyces cerevisiae. Mol Cell Biol. 20:7559–7571.

Nagiec, M.J., P.C. McCarter, J.B. Kelley, G. Dixit, T.C. Elston, and H.G. Dohlman. 2015. Signal inhibition by a dynamically regulated pool of monophosphorylated MAPK. Mol Biol Cell. 26:3359–3371.

Nishimura, A., K. Yamamoto, M. Oyama, H. Kozuka-Hata, H. Saito, and K. Tatebayashi. 2016. Scaffold Protein Ahk1, Which Associates with Hkr1, Sho1, Ste11, and Pbs2, Inhibits Cross Talk Signaling from the Hkr1 Osmosensor to the Kss1 Mitogen-Activated Protein Kinase. Mol Cell Biol. 36:1109–1123.

O’Rourke, S.M., and I. Herskowitz. 1998. The Hog1 MAPK prevents cross talk between the HOG and pheromone response MAPK pathways in Saccharomyces cerevisiae. Genes Dev. 12:2874–2886.

Park, H.O., P.J. Kang, and A.W. Rachfal. 2002. Localization of the Rsr1/Bud1 GTPase involved in selection of a proper growth site in yeast. The Journal of biological chemistry. 277:26721–26724.

Patterson, J.C., L.S. Goupil, and J. Thorner. 2021. Cdc42-Specific GTPase-Activating Protein Rga1 Squelches Crosstalk between the High-Osmolarity Glycerol (HOG) and Mating Pheromone Response MAPK Pathways. Biomolecules. 11.

Peter, M., A.M. Neiman, H.O. Park, M. van Lohuizen, and I. Herskowitz. 1996. Functional analysis of the interaction between the small GTP binding protein Cdc42 and the Ste20 protein kinase in yeast. EMBO J. 15:7046–7059.

Pitoniak, A., C.A. Chavel, J. Chow, J. Smith, D. Camara, S. Karunanithi, B. Li, K.H. Wolfe, and P.J. Cullen. 2015. Cdc42p-interacting protein Bem4p regulates the filamentous-growth mitogen-activated protein kinase pathway. Mol Cell Biol. 35:417–436.

Posas, F., and H. Saito. 1997. Osmotic activation of the HOG MAPK pathway via Ste11p MAPKKK: scaffold role of Pbs2p MAPKK. Science. 276:1702–1705.

Prabhakar, A., J. Chow, A.J. Siegel, and P.J. Cullen. 2020. Regulation of intrinsic polarity establishment by a differentiation-type MAPK pathway in S. cerevisiae. J Cell Sci. 133.

Prabhakar, A., B. González, H. Dionne, S. Basu, and P.J. Cullen. 2021. Spatiotemporal control of pathway sensors and cross-pathway feedback regulate a differentiation MAPK pathway in yeast. J Cell Sci. 134.

Pryciak, P.M., and F.A. Huntress. 1998. Membrane recruitment of the kinase cascade scaffold protein Ste5 by the Gbetagamma complex underlies activation of the yeast pheromone response pathway. Genes Dev. 12:2684–2697.

Raitt, D.C., F. Posas, and H. Saito. 2000. Yeast Cdc42 GTPase and Ste20 PAK-like kinase regulate Sho1-dependent activation of the Hog1 MAPK pathway. EMBO J. 19:4623–4631.

Raman, M., W. Chen, and M. Cobb. 2007. Differential regulation and properties of MAPKs. Oncogene. 26:3100–3112.

Ramezani Rad, M., G. Jansen, F. Bühring, and C.P. Hollenberg. 1998. Ste50p is involved in regulating filamentous growth in the yeast Saccharomyces cerevisiae and associates with Ste11p. Mol Gen Genet. 259:29–38.

Rane, C.K., and A. Minden. 2014. P21 activated kinases: structure, regulation, and functions. Small GTPases. 5.

Richman, T.J., K.A. Toenjes, S.E. Morales, K.C. Cole, B.T. Wasserman, C.M. Taylor, J.A. Koster, M.F. Whelihan, and D.I. Johnson. 2004. Analysis of cell-cycle specific localization of the Rdi1p RhoGDI and the structural determinants required for Cdc42p membrane localization and clustering at sites of polarized growth. Curr Genet. 45:339–349.

Roberts, C.J., B. Nelson, M.J. Marton, R. Stoughton, M.R. Meyer, H.A. Bennett, Y.D. He, H. Dai, W.L. Walker, T.R. Hughes, M. Tyers, C. Boone, and S.H. Friend. 2000. Signaling and circuitry of multiple MAPK pathways revealed by a matrix of global gene expression profiles. Science. 287:873–880.

Roberts, R.L., and G.R. Fink. 1994. Elements of a single MAP kinase cascade in Saccharomyces cerevisiae mediate two developmental programs in the same cell type: mating and invasive growth. Genes Dev. 8:2974–2985.

Roberts, R.L., H.U. Mösch, and G.R. Fink. 1997. 14-3-3 proteins are essential for RAS/MAPK cascade signaling during pseudohyphal development in S. cerevisiae. Cell. 89:1055–1065.

Rossman, K.L., C.J. Der, and J. Sondek. 2005. GEF means go: turning on RHO GTPases with guanine nucleotide-exchange factors. Nature reviews Molecular cell biology. 6:167.

Ryan, O., R.S. Shapiro, C.F. Kurat, D. Mayhew, A. Baryshnikova, B. Chin, Z.Y. Lin, M.J. Cox, F. Vizeacoumar, D. Cheung, S. Bahr, K. Tsui, F. Tebbji, A. Sellam, F. Istel, T. Schwarzmüller, T.B. Reynolds, K. Kuchler, D.K. Gifford, M. Whiteway, G. Giaever, C. Nislow, M. Costanzo, A.C. Gingras, R.D. Mitra, B. Andrews, G.R. Fink, L.E. Cowen, and C. Boone. 2012. Global gene deletion analysis exploring yeast filamentous growth. Science. 337:1353–1356.

Saito, H. 2010. Regulation of cross-talk in yeast MAPK signaling pathways. Current opinion in microbiology. 13:677–683.

Saito, H., and F. Posas. 2012. Response to hyperosmotic stress. Genetics. 192:289–318.

Saito, H., and K. Tatebayashi. 2004. Regulation of the osmoregulatory HOG MAPK cascade in yeast. Journal of biochemistry. 136:267–272.

Schwartz, M.A., and H.D. Madhani. 2004. Principles of MAP kinase signaling specificity in Saccharomyces cerevisiae. Annu Rev Genet. 38:725–748.

Sharmeen, N., T. Sulea, M. Whiteway, and C. Wu. 2019. The adaptor protein Ste50 directly modulates yeast MAPK signaling specificity through differential connections of its RA domain. Mol Biol Cell. 30:794–807.

Sikorski, R.S., and P. Hieter. 1989. A system of shuttle vectors and yeast host strains designed for efficient manipulation of DNA in Saccharomyces cerevisiae. Genetics. 122:19–27.

Simon, M.N., C. De Virgilio, B. Souza, J.R. Pringle, A. Abo, and S.I. Reed. 1995. Role for the Rho-family GTPase Cdc42 in yeast mating-pheromone signal pathway. Nature. 376:702–705.

Smith, G.R., S.A. Givan, P. Cullen, and G.F. Sprague, Jr. 2002. GTPase-activating proteins for Cdc42. Eukaryot Cell. 1:469–480.

Sprague, G.F., Jr., L.C. Blair, and J. Thorner. 1983. Cell interactions and regulation of cell type in the yeast Saccharomyces cerevisiae. Annu Rev Microbiol. 37:623–660.

Stevenson, B.J., B. Ferguson, C. De Virgilio, E. Bi, J.R. Pringle, G. Ammerer, and G.F. Sprague, Jr. 1995. Mutation of RGA1, which encodes a putative GTPase-activating protein for the polarity-establishment protein Cdc42p, activates the pheromone-response pathway in the yeast Saccharomyces cerevisiae. Genes Dev. 9:2949–2963.

Stevenson, B.J., N. Rhodes, B. Errede, and G.F. Sprague, Jr. 1992. Constitutive mutants of the protein kinase STE11 activate the yeast pheromone response pathway in the absence of the G protein. Genes Dev. 6:1293–1304.

Tanaka, K., K. Tatebayashi, A. Nishimura, K. Yamamoto, H.-Y. Yang, and H. Saito. 2014. Yeast osmosensors Hkr1 and Msb2 activate the Hog1 MAPK cascade by different mechanisms. Sci. Signal. 7:ra21–ra21.

Tatebayashi, K., K. Tanaka, H.Y. Yang, K. Yamamoto, Y. Matsushita, T. Tomida, M. Imai, and H. Saito. 2007. Transmembrane mucins Hkr1 and Msb2 are putative osmosensors in the SHO1 branch of yeast HOG pathway. The EMBO journal. 26:3521–3533.

Tatebayashi, K., K. Yamamoto, M. Nagoya, T. Takayama, A. Nishimura, M. Sakurai, T. Momma, and H. Saito. 2015. Osmosensing and scaffolding functions of the oligomeric four-transmembrane domain osmosensor Sho1. Nature Communications. 6:6975.

Tatebayashi, K., K. Yamamoto, K. Tanaka, T. Tomida, T. Maruoka, E. Kasukawa, and H. Saito. 2006. Adaptor functions of Cdc42, Ste50, and Sho1 in the yeast osmoregulatory HOG MAPK pathway. EMBO J. 25:3033–3044.

Tong, Z., X.D. Gao, A.S. Howell, I. Bose, D.J. Lew, and E. Bi. 2007. Adjacent positioning of cellular structures enabled by a Cdc42 GTPase-activating protein-mediated zone of inhibition. J Cell Biol. 179:1375–1384.

Truckses, D.M., J.E. Bloomekatz, and J. Thorner. 2006. The RA domain of Ste50 adaptor protein is required for delivery of Ste11 to the plasma membrane in the filamentous growth signaling pathway of the yeast Saccharomyces cerevisiae. Mol Cell Biol. 26:912–928.

Vadaie, N., H. Dionne, D.S. Akajagbor, S.R. Nickerson, D.J. Krysan, and P.J. Cullen. 2008. Cleavage of the signaling mucin Msb2 by the aspartyl protease Yps1 is required for MAPK activation in yeast. J Cell Biol. 181:1073–1081.

van der Felden, J., S. Weisser, S. Brückner, P. Lenz, and H.U. Mösch. 2014. The transcription factors Tec1 and Ste12 interact with coregulators Msa1 and Msa2 to activate adhesion and multicellular development. Mol Cell Biol. 34:2283–2293.

Van Drogen, F., N. Dard, S. Pelet, S.S. Lee, R. Mishra, N. Srejić, and M. Peter. 2020. Crosstalk and spatiotemporal regulation between stress-induced MAP kinase pathways and pheromone signaling in budding yeast. Cell Cycle. 19:1707–1715.

van Hemert, M.J., G.P. van Heusden, and H.Y. Steensma. 2001. Yeast 14-3-3 proteins. Yeast. 18:889–895.

van Heusden, G.P., D.J. Griffiths, J.C. Ford, A.W.T.F. Chin, P.A. Schrader, A.M. Carr, and H.Y. Steensma. 1995. The 14-3-3 proteins encoded by the BMH1 and BMH2 genes are essential in the yeast Saccharomyces cerevisiae and can be replaced by a plant homologue. Eur J Biochem. 229:45–53.

van Heusden, G.P., and H.Y. Steensma. 2006. Yeast 14-3-3 proteins. Yeast. 23:159–171.

Vázquez-Ibarra, A., G. Rodríguez-Martínez, G. Guerrero-Serrano, L. Kawasaki, L. Ongay-Larios, and R. Coria. 2020. Negative feedback-loop mechanisms regulating HOG- and pheromone-MAPK signaling in yeast. Curr Genet. 66:867–880.

Wai, S.C., S.A. Gerber, S.P. Gygi, and R. Li. 2007. Multisite phosphorylation of the exchange factor Cdc24 during yeast cell polarization. Regulation of the Rho GTPase Cdc42 During Cell Polarization in Budding Yeast.

Wai, S.C., S.A. Gerber, and R. Li. 2009. Multisite phosphorylation of the guanine nucleotide exchange factor Cdc24 during yeast cell polarization. PloS one. 4:e6563.

Weisz Hubsman, M., N. Volinsky, E. Manser, D. Yablonski, and A. Aronheim. 2007. Autophosphorylation-dependent degradation of Pak1, triggered by the Rho-family GTPase, Chp. Biochem J. 404:487–497.

Winters, M.J., and P.M. Pryciak. 2005. Interaction with the SH3 domain protein Bem1 regulates signaling by the Saccharomyces cerevisiae p21-activated kinase Ste20. Molecular and cellular biology. 25:2177–2190.

Winzeler, E.A., D.D. Shoemaker, A. Astromoff, H. Liang, K. Anderson, B. Andre, R. Bangham, R. Benito, J.D. Boeke, H. Bussey, A.M. Chu, C. Connelly, K. Davis, F. Dietrich, S.W. Dow, M. El Bakkoury, F. Foury, S.H. Friend, E. Gentalen, G. Giaever, J.H. Hegemann, T. Jones, M. Laub, H. Liao, N. Liebundguth, D.J. Lockhart, A. Lucau-Danila, M. Lussier, N. M’Rabet, P. Menard, M. Mittmann, C. Pai, C. Rebischung, J.L. Revuelta, L. Riles, C.J. Roberts, P. Ross-MacDonald, B. Scherens, M. Snyder, S. Sookhai-Mahadeo, R.K. Storms, S. Véronneau, M. Voet, G. Volckaert, T.R. Ward, R. Wysocki, G.S. Yen, K. Yu, K. Zimmermann, P. Philippsen, M. Johnston, and R.W. Davis. 1999. Functional characterization of the S. cerevisiae genome by gene deletion and parallel analysis. Science. 285:901–906.

Wu, C., G. Jansen, J. Zhang, D.Y. Thomas, and M. Whiteway. 2006. Adaptor protein Ste50p links the Ste11p MEKK to the HOG pathway through plasma membrane association. Genes & development. 20:734–746.

Xu, G., G. Jansen, D.Y. Thomas, C.P. Hollenberg, and M. Ramezani Rad. 1996. Ste50p sustains mating pheromone-induced signal transduction in the yeast Saccharomyces cerevisiae. Mol Microbiol. 20:773–783.

Yamamoto, K., K. Tatebayashi, K. Tanaka, and H. Saito. 2010. Dynamic control of yeast MAP kinase network by induced association and dissociation between the Ste50 scaffold and the Opy2 membrane anchor. Mol Cell. 40:87–98.

Yang, Z., H.M. Kirton, D.A. MacDougall, J.P. Boyle, J. Deuchars, B. Frater, S. Ponnambalam, M.E. Hardy, E. White, S.C. Calaghan, C. Peers, and D.S. Steele. 2015. The Golgi apparatus is a functionally distinct Ca2+ store regulated by the PKA and Epac branches of the β1-adrenergic signaling pathway. Sci Signal. 8:ra101.

