## Supplementary figures and images for "New Features Surrounding the Cdc42-Ste20 Module that Regulates MAP Kinase Signaling in Yeast"

### Fig. S1

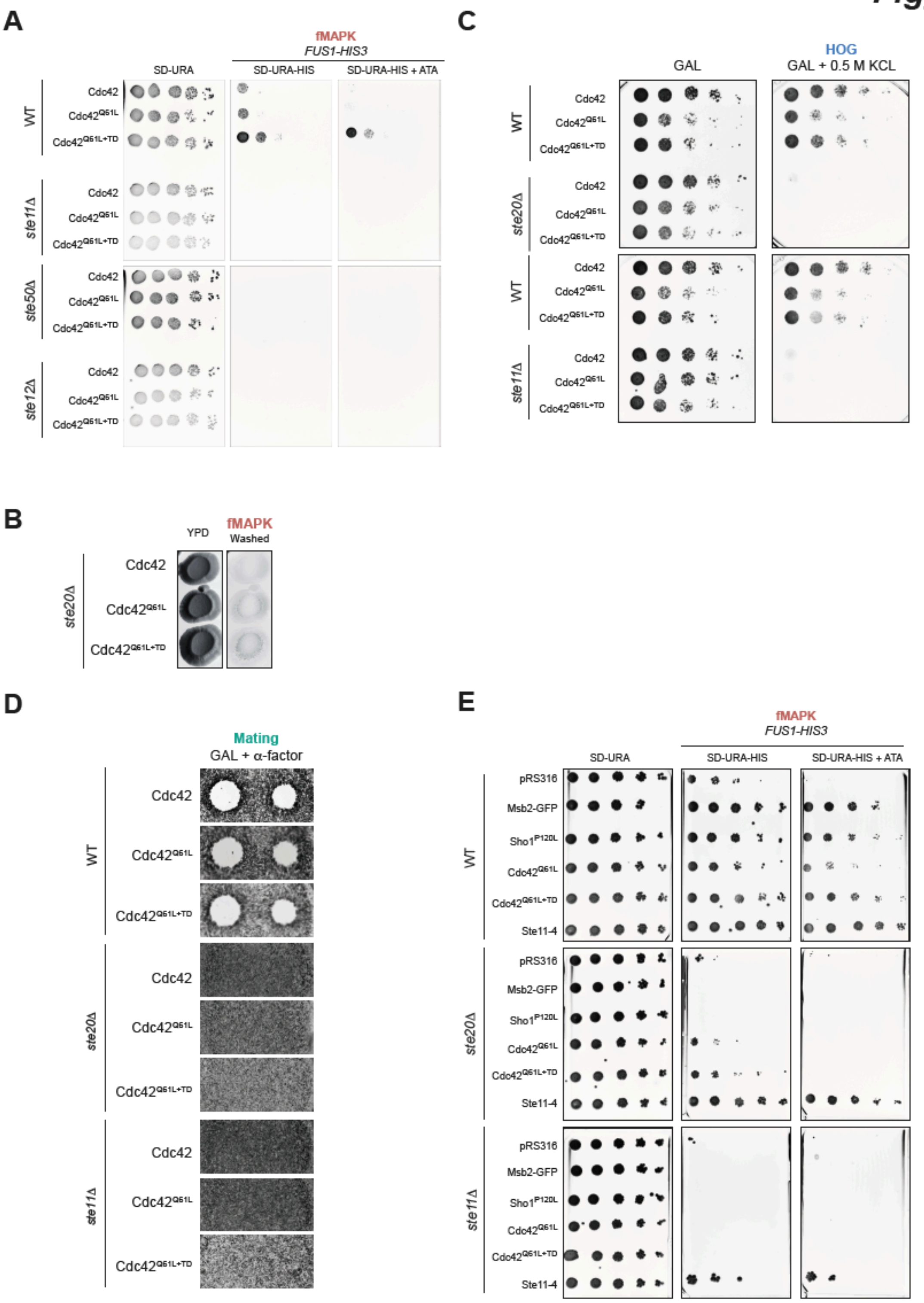

### Fig. S2

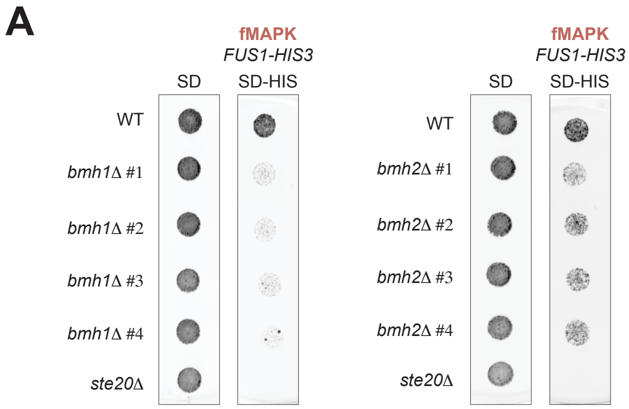

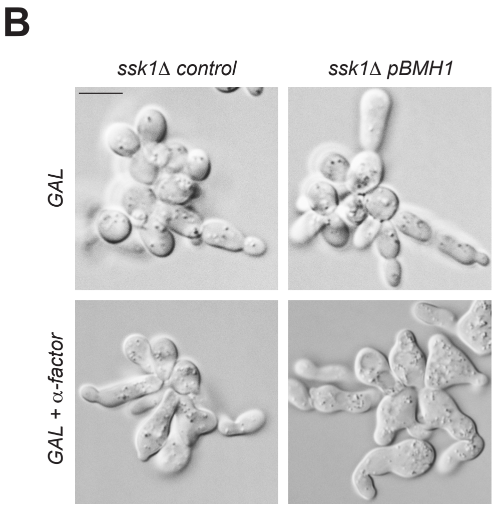

### Fig. S3

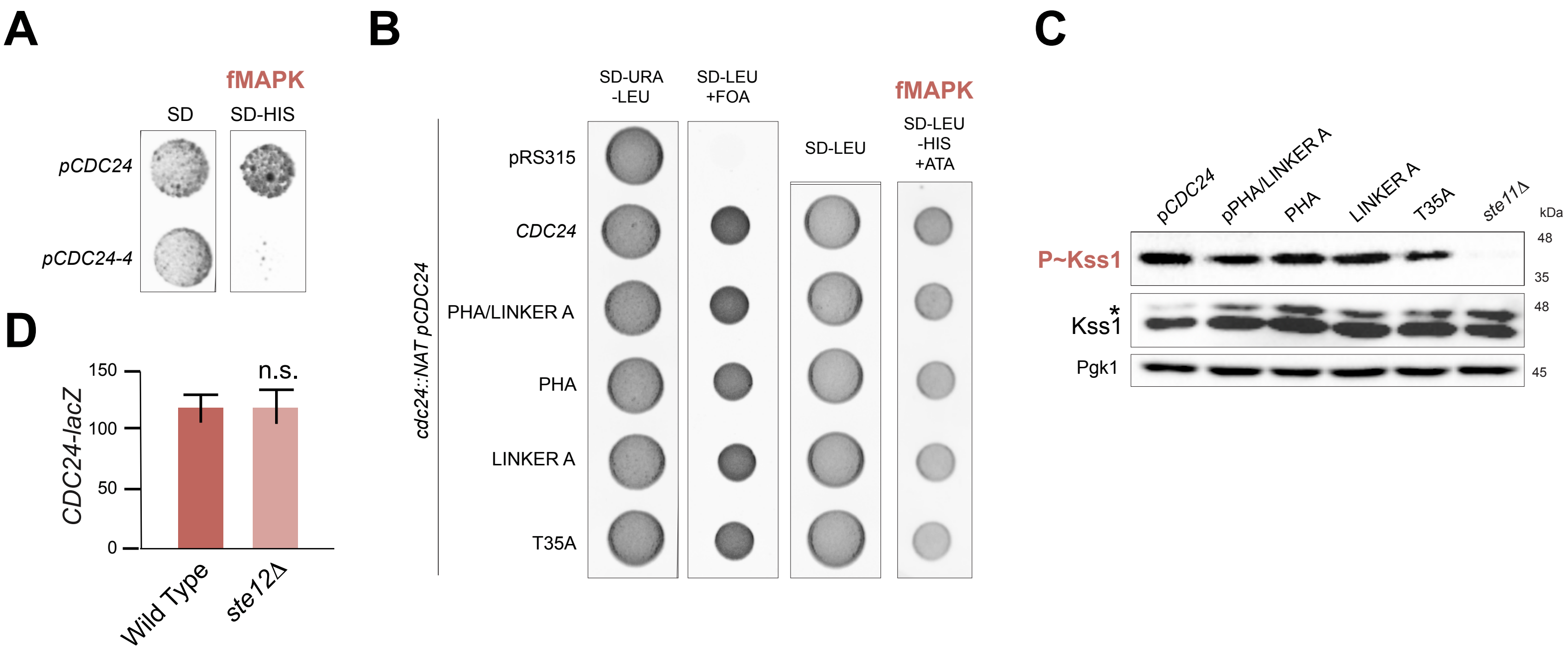
